# Encoding of antennal position and velocity by the Johnston’s organ in hawkmoths

**DOI:** 10.1101/2024.07.28.605492

**Authors:** Chinmayee L Mukunda, Sanjay P Sane

## Abstract

Insect antennae function as versatile, multimodal sensory probes in diverse behavioural contexts. In addition to their primary role as olfactory organs, they serve essential mechanosensory functions across insects, including auditory perception, vestibular feedback, airflow detection, gravity sensing, and tactile sensation. These diverse functions are facilitated by the mechanosensory Johnston’s organ (JO), located at the joint between the second antennal segment, known as the pedicel, and the flagellum. The pedicel-flagellum joint lacks muscles which means that the Johnston’s organs can perceive only passive deflections of the flagellum. Earlier work which characterized the sensitivity and short response time of the sensory units of JO in hawkmoths, showed that their sensitivity to a broad frequency range is range-fractionated. This vastly expands the functional repertoire of the JO. However, it is not clear what components of antennal kinematics are encoded by the JO. Here, we conducted experiments to test the hypothesis that JO neurons encode the position and velocity of angular movements of the flagellum. We recorded intracellularly from the axons of primary sensory neurons of JO while stimulating it with ramp-and-hold stimuli in which antennal position or antennal angular velocity was maintained at various constant values. Our study shows that JO neurons encode angular velocity and position of the antenna in their response. We also characterized the neural adaptation of the responses to angular velocities and positions. A majority of neurons were sensitive to a movement in the ventrad direction, in the direction of gravity. The adaptation and the directional response properties give rise to a nonlinear hysteresis-like response. Together, these findings highlight the neurophysiological basis underlying the functional versatility of the JO.

## INTRODUCTION

Despite the staggering diversity in forms and shapes, the insect antenna is a remarkably conserved multisensory organ. It is involved in multimodal functions including olfaction, thermoception, hygroception, and mechanosensation. The mechanosensors on the insect antennae mediate various important functions including tactile sensing (in cockroaches; (Camhi and Johnson, 1999; Okada and Toh, 2000; Comer and Baba, 2011), audition (in mosquitos, honeybees, and flies; Johnston, 1855; Dreller and Kirchner, 1993; Eberl, 1999), and vestibular feedback (in Lepidoptera; Niehaus, 1981; Sane *et al*., 2007; Sane, Manjunath and Mukunda, 2023). The antennae of all Neopteran insects consist of three segments: a basal scape, a middle pedicel, and a distal flagellum (Schneider, 1964; Krishnan *et al*., 2012). Whereas the head-scape and scape-pedicel movements are actuated by the extrinsic and intrinsic muscles respectively, movements of the flagellum relative to the pedicel are passive and moved only by external perturbations or indirectly due to body movements (Kloppenburg *et al*., 1997; Sant and Sane, 2018). These passive flagellar deflections are detected by an extremely sensitive mechanosensory chordotonal organ, the Johnston’s organ (JO), which spans across the pedicel-flagellum joint (Johnston, 1855; Gewecke, 1974). Johnston’s organs serve diverse functions across insects. For instance, honeybee JOs are thought to perceive the auditory cues generated by a forager bee during waggle dances (Dreller and Kirchner, 1993; Dreller and Kirchner, 1993) as well as airflow during flight (Heran, 1957; Khurana and Sane, 2016). In mosquitoes, JO neurons encode the auditory stimuli generated during flight and courtship (Gopfert and Robert, 2001; Lapshin and Vorontsov, 2017). These mechanosensors are also involved in antennal positioning and providing flight-related feedback in blowflies, locusts, hawkmoths, and vinegar flies (Burkhardt and Gewecke, 1965; Gewecke, 1967; Sane *et al*., 2007; Mamiya and Dickinson, 2015; Suver *et al*., 2019) and tactile substrate vibration sensing in stink bugs (Jeram and Čokl, 1996). In hawkmoths, JOs provide vestibular feedback during flight (Sane et al 2007; Dahake et al, 2018; Sane et al 2023), airflow-dependent antennal positioning (Khurana and Sane, 2016; Natesan *et al*., 2019), head stabilization (Chatterjee *et al*., 2022), and flower-tracking behaviours (Dahake *et al*., 2018). Together, this body of work demonstrates both the range and diversity of JO function. In all the above cases, JO is activated *via* the movement of the flagellum relative to the pedicel and hence its basic mechanism is conserved despite the diversity.

The Johnston’s organ is composed of several sensory units called scolopidia. Each scolopidial unit consists of dendrites of 2-3 sensory neurons that project to a scolopale cell which connects to the cuticular invaginations of the flagellum (Field and Matheson, 1998). Mechanoreceptors at the dendritic tip transduce the mechanical vibrations of the flagellum into receptor potentials (Zanini and Göpfert, 2014; Eberl, Kamikouchi and Albert, 2016). The cell bodies of the sensory neurons located in the pedicel integrate these receptor potentials and generate action potentials that encode the movements of the flagellum (Kamikouchi *et al*., 2009). This encoded information can be used by the nervous system as sensory feedback to modulate behaviours.

To perceive subtle perturbations of the flagella during flight manoeuvres, the sensory neurons need to be sensitive to extremely small deflections and must respond very rapidly at high frequencies. Intracellular electrophysiological recordings in hawkmoth JO confirmed that sensory neurons respond to deflections of as little as ∼0.05 degrees at latencies as low as 3 ms (Dieudonne, Daniel and Sane, 2014). JO is sensitive to a wide range of frequencies and range fractionated, which means that individual scolopidia do not trade off sensitivity and range, but are individually finely tuned to a narrow frequency range (in hawkmoths *Manduca sexta*-Sane *et al*., 2007; Dieudonne, Daniel and Sane, 2014; in mosquitoes *Culex pipiens* Lapshin and Vorontsov, 2017). Similar characteristics have also been observed through calcium imaging of JO neuron activity in *Drosophila* (Kamikouchi *et al*., 2009; Yorozu *et al*., 2009; Patella and Wilson, 2018).

Because mechanosensory neurons encode diverse mechanical stimuli across insects, the understanding of the fundamental features of the stimulus encoded by these neurons is crucial to determine the mechanisms that underlie the function. Specifically, what components of the ambient, analog mechanical stimuli are encoded by JO neurons? Previous investigations of this question have used diverse stimuli including sinusoids, chirps, and band-limited Gaussian noise stimuli to characterize response to a wide range of stimulus frequencies (Sane *et al*., 2007; Dieudonne, Daniel and Sane, 2014). These studies shed light on specific encoding properties of the JO neurons in relation to their function as antennal mechanosensors. However, an inherent limitation in the stimulus delivery system in these studies meant that the stimuli were mechanically-filtered and hence, a readout of the delivered stimulus closest to the sensor of interest is essential for accurate interpretation of the results of these experiments. Moreover, white noise stimuli scramble from the stimulus all of its natural context, which may render the resulting responses to be very difficult to interpret. Indeed, this lacuna is typical of any “systems level” white noise characterization of sensory encoding properties.

An alternative approach involves examining the system’s response to individual components of a complex stimulus. This method aims to gain insights from the system’s reactions to simpler stimuli, which can then aid in understanding its responses to more complex stimuli. Using this latter approach, we presented stimuli designed to address the encoding properties of JO neurons. Specifically, we proposed that the neural activity of JO neurons encodes the angular position and velocity of flagellar movement, which represent the initial terms in the series expansion of any complex stimulus time series function. We presented the antenna of the Oleander hawkmoth *Daphnis nerii* with either constant position or velocity stimuli, while simultaneously performing intracellular recordings from axons of JO neurons in the antennal nerve. Here we describe the encoding properties of JO neurons to components of a movement stimulus. We also discuss the similarities observed in the physiology of the JOs with femoral chordotonal organs and campaniform sensilla in the legs of insects, suggesting common underlying encoding principles across these mechanosensors.

## MATERIALS AND METHODS

### Moth culture

Wild *Daphnis nerii* larvae or pupae initially collected from the surrounding area were bred in a large cage housing host plants and artificial nectaries. The female moths lay eggs on the leaves of *Nerium oleander* or *Tabernaemontana divaricata* plants after mating. The eggs were collected in plastic containers. The hatched larvae were provided with tender Nerium leaves. The larvae were transferred into larger boxes with harder leaves as they grew old. *Daphnis* undergoes 5 molts or “instars” in the larval phase. Fifth instar larvae were kept in sawdust for pupation which provided sufficient friction to remove their pupal case during emergence. The pupal stage lasts for ∼15 days, after which the late-stage pupae were transferred into an emergence cage from where the moths were collected for experiments.

### Preparation

Adult oleander hawkmoths *Daphnis nerii* were used for all the experiments. Healthy male or female moths were collected from the moth culture facility 0–1 day post-eclosion. The moths were visually inspected for any deformities in the body and for normal activity levels to assess that they were healthy and active.

*Immobilization.* The head, neck, thorax, and wing bases were descaled using a toothbrush for proper immobilization. The legs were detached at the coxa-trochanter joint, to minimize haemolymph loss. The wings, thoracic sclerites, thorax-abdomen joint, and the head-neck joint were immobilized using hot dental wax. The moth was placed dorsal side up in a customized plastic syringe tube and attached to the tube using hot wax; the dorsal side of the head was kept accessible for antennal stimulation and surgery (**Fig 1A i**).

**Figure 1.**
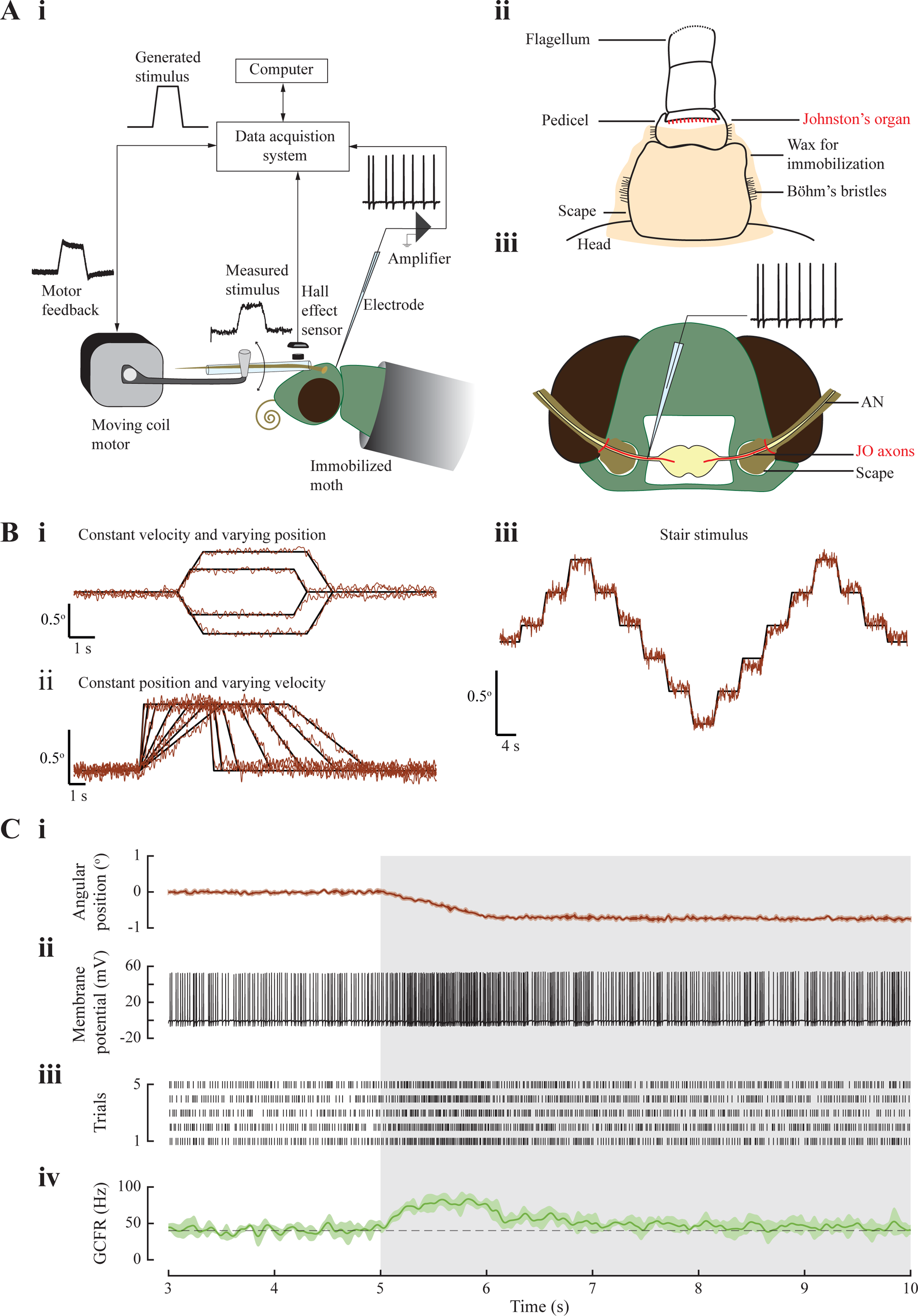
Experimental setup and processing of raw data. (A i) Schematic of the experimental setup for intracellular recording from axons of single neuronal units of the antennal mechanosensory Johnston’s organ. A custom-made capillary attachment connects the flagellum to the motor. A Hall effect sensor placed opposed to a neodymium magnet mounted on the capillary records the actual movement delivered. (A ii) The head-scape and scape-pedicel joints of the antenna are immobilized using a mixture of beeswax and Rosin (3:2). This ensures that the stimulus acts only on the pedicel-flagellum joint where the Johnston’s organ is located. (A iii) Schematic of the surgery performed to expose the moth brain and the antennal nerve. The axons of JO (in red) neurons project to the brain through the antennal nerve. The recording electrode is inserted into the antennal nerve (AN). (B) Comparison of computer-generated stimulus and the filtered output of the Hall sensor. Ramp-and-hold stimuli with constant angular velocity and variable angular positions (B i), constant angular position and variable angular velocities (B ii), constant velocity and constant relative change in position – stair protocol (B iii). (C) Analysis pipeline showing the processing of raw data. (i) Ventrad ramp-and-hold stimulus delivered to the flagellum, as measured by the Hall effect sensor. *Solid line*: mean of 5 trials, *shaded region*: 1 Standard Deviation. The grey box marks the stimulus duration following a rest period; (ii) representative trace showing single trial of recorded membrane potential in response to the above stimulus. Action potentials (or spikes) with hyperpolarization are indicative of intracellular recording; (iii) raster plot showing spike timing in 5 trials of the same stimulus; (iv) Gaussian Convolved Firing Rate (GCFR) is a smoothened histogram of firing rate (mean ± SD, *solid line* and *shaded region* respectively) and dashed line marks the average baseline firing rate.

*Positioning antenna for stimulus.* The tube holding the moth was mounted on a stand in the electrophysiology rig at an angle of ∼40°. The flagellum of the left antenna was passed through a custom-made capillary, which was attached to the motor shaft. The antenna was positioned at ∼45° to the longitudinal axis of the moth. The head-scape and scape-pedicel joints of the left antenna were immobilized (**Fig 1A ii**) by depositing a mixture of beeswax and rosin (3:2), using a custom-made cauterizer with fine filament (∼0.5 mm tip). The two joints were immobilized to specifically stimulate JO and restrict the activity of the antennal musculature. The left flagellum was connected to the capillary attachment at the 3^rd^ annulus of the flagellum. All the joints of the right antenna were immobilized with wax to minimize movements in the preparation.

*Surgery.* The cuticle of the head and the tissue underneath were cut out to expose the brain and the antennal nerve (**Fig 1A iii**). The trachea on the nerve were removed using fine forceps for easy penetration of the nerve by the electrode. The antennal nerve was supported from underneath by a bent syringe needle (PrecisionGlide^®^ Needle, 305128). The tip of the needle was coated with a resin-based adhesive (Araldite^®^) and dried, thus eliminating slippage of the nerve and provided stability for recordings. This needle also delivered insect saline (recipe below) to the preparation.

### Electrophysiology

Intracellular recordings were performed on individual axons of the JO neurons passing through the antennal nerve. The exposed region was constantly perfused with physiological insect saline. The saline consisted of 150 mmol l^-1^ NaCl, 3 mmol l^-1^ CaCl2, 3 mmol l^-1^ KCl, 10 mmol l^-1^ N-tris (hydroxymethyl) methyl-2-aminoethane sulfonic acid buffer and 25 mmol l^-1^ sucrose, pH 6.5–7.5 (Lei, Christensen and Hildebrand, 2004; Dieudonne, Daniel and Sane, 2014). Chloridized silver wires were used as recording and reference electrodes. The reference electrode was placed in the vicinity of the antennal nerve and dipped in the saline perfusion. A quartz capillary (outer diameter: 1 mm, inner diameter: 0.5 mm, with filament, Sutter Instrument, Novato, CA 94949, USA) was pulled to standardized dimensions using a customized program in the pipette puller (P-2000, Sutter Instrument, Novato, CA 94949, USA). The recording electrode was placed in the pulled pipette filled with 1M KCl.

The recording electrode was connected to the amplifier head stage through a 3D-printed custom electrode holder. The head stage was mounted on a micromanipulator (EMM2, Narishige, Japan) that controlled the precise position of the electrode. The recorded activity was amplified with a gain of 10 (IX2 pre-amplifier, Dagan Corporation, Minneapolis, Minnesota, United States). All the signals were registered in the computer through the data acquisition system (USB-6363, NI, Austin, Texas, USA) (Fig 1Ai).

### Stimulus delivery

The capillary enclosing the flagellum was connected to a motor (300C-I, Aurora scientific, Canada) through a linkage system (**Fig 1A**). This system was driven by computer commands delivered through the output channel of the data acquisition system (USB-6363, NI, Austin, Texas, USA). The stimulus protocols were generated by a custom-written MATLAB program (The MathWorks, Inc. (2023), Natick, Massachusetts, United States).

The actual stimulus experienced by the antenna is considerably different from the intended stimulus, as it is altered by the mechanical apparatus linking the motor to the antenna. It is thus necessary to measure the actual stimulus, especially when working with highly-sensitive mechanosensors. The stimuli delivered in all our experiments were along a single axis (dorsoventral). To obtain a readout of the actual stimulus delivered to the JO, a neodymium magnet (diameter = 3 mm, thickness = 2 mm) was mounted close to the tip of the capillary holding the flagellum. A Hall-effect sensor aligned with the magnet was used to measure the change in the magnetic field as the flagellum moved upon stimulus delivery. The Hall-effect sensor thus provided a more faithful readout of the actual stimulus delivered, closer to the base of the antenna. Although the values mentioned below also indicate the intended stimuli (angular displacements and angular velocities), all the analyses and related plots were obtained using the actual readouts from the Hall-effect sensor as stimuli. A voltage versus distance calibration curve obtained for the sensor-magnet pair provided the actual distances by which the capillary moved. The distance between the 3^rd^ annulus and the pedicel-flagellum joint was measured in every moth at the end of the experiment session under a microscope (SMZ25, Tokyo, Japan) with calibrated scale in the associated software (NIS-Elements). The actual angular movements delivered to the JO were calculated using the formula:

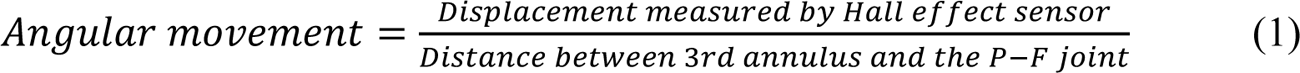

### Stimulus protocols

The MATLAB-generated stimulus protocols were designed to obtain both the static and dynamic responses of the neurons (**Fig 1B i-iii)**. A step stimulus has been typically used to estimate the response latencies, adaptation dynamics, and encoding properties of mechanosensors (Chapman and Smith, 1963; Dieudonne, Daniel and Sane, 2014). An ideal step stimulus involves an instantaneous change in position, which is not possible for a mechanical system to achieve. The neuron may respond to the sudden deflection in position, velocity or acceleration of the movement, making it difficult to identify and delineate which components of the delivered stimulus elicited the response. To overcome this issue, we used ramp-and-hold stimuli in which angular velocities and angular positions could be independently varied. This allowed us to decouple the response of neurons to angular velocities and positions. Similar stimulus protocols have been used in the physiological characterization of femoral chordotonal organs (Hofmann, Koch and Bässler, 1985), campaniform sensilla (Ridgel *et al*., 2000) and mechanosensory descending neurons (Ache and Dürr, 2013).

In the experiments described here, the range of hold positions tested were limited by how much the flagellum could be moved without affecting the stability of the recording. Due to limitations of the motor, there was also a trade-off between the amplitude and velocity of movement. Hence, it was not possible to attain the same hold positions exceeding 2 orders of magnitude change in constant angular velocity.

The protocols were as follows:

*Ramp-and-hold stimuli with variable hold positions.* The flagellum was moved to different positions (1°or 0.5° above and below baseline position) at a fixed angular velocity of 1°/second (**Fig 1B i**).

*Ramp-and-hold stimuli with variable ramp velocities.* The flagellum moved by 1° at different velocities (0.1 to 4 °/s) and held at constant position for 4 seconds (**Fig 1B ii**).

*Stair.* A stair protocol spanned 2° in steps of 0.4°. The transitions between the positions were made at a constant velocity of 0.04°/s in both dorsad and ventrad directions. Each step lasted for 4 seconds (**Fig 1B iii**).

A resting period of 6-10 seconds (corresponding to approximately 180-300 wing beats) was provided between consecutive trials, allowing the neural activity to return to the baseline level. Positive and negative values of angular position or velocity correspond to a dorsad (towards dorsal side of the moth) or a ventrad (towards ventral side) movement of the flagellum, respectively. The effective values of angular positions and velocities depended on the point of attachment of the flagellum to the capillary which varied across preparations.

Nevertheless, the stimulus presentations were reliably repeatable within recordings from individual moths. For example, if the actual deflection was 0.8° when the target deflection was 1°, this value was consistently achieved across all trials and protocols tested on that moth. The angular velocity profile of the stimulus signal is displayed in the middle panels of **Fig 2A**, 3A (also **Supplementary Fig 1**) The velocity signal was obtained by taking the derivative of the position signal from the Hall effect sensor and filtering with a lowpass Butterworth filter (order = 3, cut-off = 4 Hz). This filtering causes the peaks to be attenuated. Thus, the reported values of velocities do not match the peak of the velocity signals in these plots. To get a reliable estimate of the velocity of the ramp, we fit a straight line to the ramp. The r^2^ of these fits was always greater than 0.9. The slope of the fit is the reported constant angular velocity value during the ramp stimulus.

**Figure 2.**
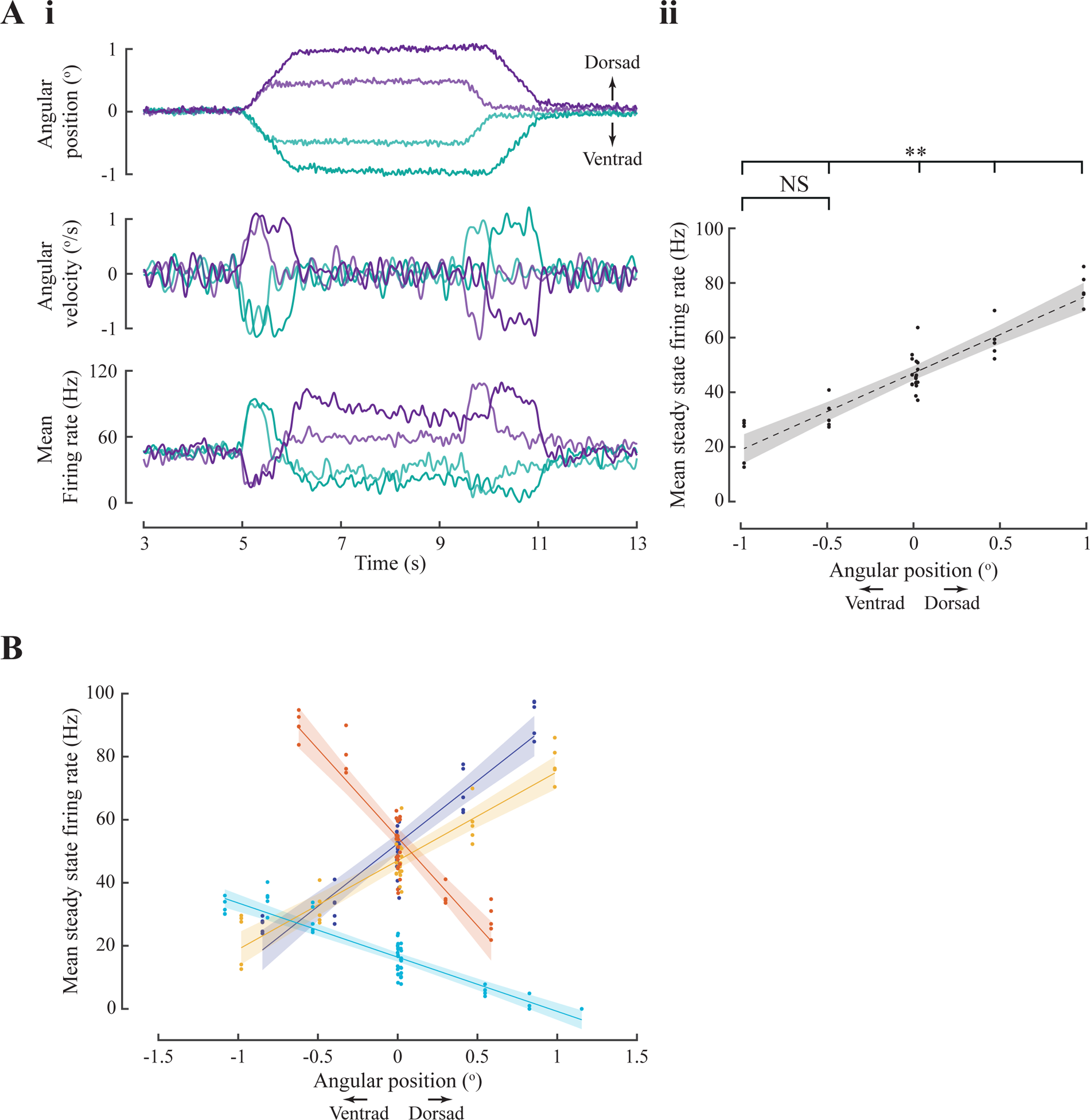
Position encoding neurons. (A) A neuron sensitive to both angular position and velocity. (A i) (*top*) Mean angular position stimulus delivered to flagellum (5 trials per protocol). The flagellum was deflected by ca. 0.5° and 1° above and below baseline. (*middle*) Angular velocity profile of the stimulus shows the same magnitude of velocity during the ramp phases to both positions in the same direction. (*bottom*) Average firing rate obtained from 5 trials for each protocol. (A ii) Mean steady-state firing rate is measured as a mean firing rate over 2.5 s to 3.5 s of the constant position stimulus. The steady-state firing rate was significantly different for all positions (Wilcoxon rank sum test, p < 0.01) except for 0.5° and 1° below baseline (p > 0.05). The steady-state firing rate linearly increased with angular position. *Dashed line:* linear fit, *shaded region:* 95% confidence interval. (B) Steady-state firing rate response to constant angular positions in 4 out of 20 neurons that showed linear relation with position with r^2^ ≥ 0.8.

### Data Analysis

For all the analyses presented here, we have used only recordings with at least 3 trials per protocol. All the data analyses were performed in MATLAB.

*Spike detection.* As described above, the prescribed stimulus was transformed by the mechanical system, and hence we used a Hall-effect sensor to record the actual stimulus close to the antennal base (**Fig 1 C i**). The recorded intracellular response was bandpass-filtered with a Butterworth filter (order = 8, cut-off = 10-2000 Hz) to eliminate drifts, baseline offsets, and high-frequency noise. 30% of the maximum amplitude of membrane voltage within a trial was set as the threshold for spike detection (**Fig 1C ii**). The spike timings across the trials were collated to generate a raster (**Fig 1C iii**). Trials with membrane voltage lower than 5 mV were discarded, as the small amplitude spikes may be indistinguishable from extracellular spikes from adjacent neurons.

*Gaussian convolved firing rate (GCFR).* A smoothened firing rate was obtained by convolving the raster data with a Gaussian kernel of 200 ms length and 30 ms width (**Fig 1C iv**).

*Baseline firing rate.* Firing rate before the start and end of a stimulus protocol. Most neurons showed some baseline activity during the resting periods while the flagellum is held at a constant angle. For instance, the neuron shown in **Figure 1C** had a mean baseline firing rate of 40 Hz which increased to a maximum of 83 Hz, when the antenna was moved ventrally and then fell to 46 Hz when the antenna was again at a new steady position of 0.7° (Fig 1C iv).

*Definitions:* The following definitions apply to the recordings and analysis in this paper.

*Mean baseline firing rate.* The mean firing rate calculated over 1 s duration of the trial-averaged firing rate, preceding the onset of the ramp stimulus by 0.5 s.

*Peak firing rate.* Maximum firing rate attained within the duration of the ramp stimulus.

*Steady-state firing rate.* Firing rate attained after the adaptation to constant position stimulus.

*Mean steady-state firing rate.* The mean firing rate calculated over 1 s duration of the trial-averaged firing rate, preceding the end of hold stimulus by 0.5 s.

*Statistical tests.* We used the Wilcoxon rank sum test to compare mean steady-state firing rates in response to different angular positions, unless mentioned otherwise. As there were 5 trials per protocol, we did not use a parametric test. A p-value of 0.05 and lower was considered significant. Neurons with significant difference in steady-state firing rates are considered to be position-sensitive.

*Quantifying the relation between the constant angular velocity stimulus (ramp) and corresponding peak firing rate*

To understand how peak firing rate relates to angular velocity, we tested 4 models from the literature and compared how well each one explained the data. Details of these 4 models are given in the **Supplementary figure 2** along with a table of fit parameters (**Supplementary Table 1**). In 3 out of 30 neurons, the response to ramp stimulus was not distinguishable from baseline. These were excluded from this analysis.

Various studies in the past have used power law and semilog fits to model the relation between firing rate and the first-order derivative of a stimulus. Technically, 2 orders of magnitude change in stimulus parameter and firing rate are required to establish power law relation. Although these models fit our data well, because the sampled velocity range was less than 2 decades, we could not justify using either power law or semilog fits. Of the 4 models, the logistical model provided the best fit (r^2^ ≥ 0.9) for most neurons, and hence this was adopted for all analysis.

*Logistic fit.* Some neurons showed a saturating nonlinear response to angular velocity. We modelled the nonlinear relation as a logistic (sigmoid) equation (Landolfa and Miller, 1995)

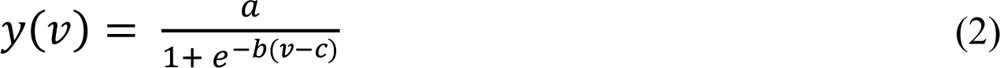

where,

*y*(*v*) = Peak firing rate, *v* = Angular velocity of flagellar movement,

*a*, *b*, *c* = Coefficients of the equation.

*Quantifying adaptation.* The ramp-and-hold stimulus is composed of a constant velocity and a constant position segment. To study the adaptation to each of these stimuli, we separately extracted the responses to the two stimuli. We quantified the adaptation to constant angular velocity (or ramp) stimulus by extracting the response from 90% of the maximum firing rate to the end of the ramp stimulus time point. To quantify adaptation to the hold stimulus, the response 100 ms after the end of the ramp stimulus to 500 ms before the end of the hold stimulus was considered. Single decaying exponential fits have been commonly used to model adaptation in neurons. (e.g. Hildebrandt, Benda and Hennig, 2009; Clemens, Ozeri-Engelhard and Murthy, 2018). Here, we modelled the adaptation of the firing rate with two separate single exponential fits.

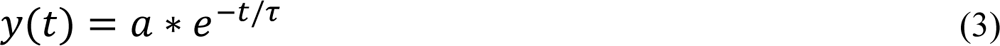

Where, *y*(*t*) = Firing rate, *t* = time, *τ* = Adaptation time constant, *a* = Scaling coefficient

*τ* < 0 implies adaptation.

*τ* = 0 implies no adaptation.

*τ* > 0 implies an increasing response.

In many cases, the r^2^ values were low due to fluctuations in the firing rate. Only fits with r^2^ ≥ 0.75 were considered for further analysis.

## RESULTS

The data from all the recorded neurons (60 neurons, from 32 moths) and their characteristics is summarized in **Table 1**. Neuronal recordings used as specific case examples are highlighted wherever relevant (**Supplementary table 2**). Most neurons showed some baseline activity during the resting period when the flagellum was held in a constant position, indicating the presence of residual deformation in the joint. In 56 out of 60 neurons, the firing rate increased relative to the baseline when the flagellum was moved in the ventrad direction. In the remaining 4 neurons, the response was ambiguous. The response of the remaining 3 neurons to movement in either direction was noisy, and therefore excluded from further analysis. To understand how the neuronal response encodes angular positions and velocities, we used ramp-and-hold stimuli in which either the hold position (i.e. angular position) was varied at constant ramp slope, or the ramp slope (i.e. angular velocity) was varied at constant hold positions. Below we describe activity of neurons encoding angular position, angular velocity, as well as exhibiting adaptation, history-dependence and directional sensitivity.

### JO neurons encoding angular position of the flagellum

We moved the flagellum along a ramp-and-hold pattern to different positions at the same velocity (**Fig 2A i, top and middle panels**). This stimulus was presented to 20 neurons of which 18 exhibited an initial rise or fall in firing rate during the ramp phase, depending on the direction of the ramp. This was followed by an easing (or adaptation) in firing rate during the hold phase, or sometimes even prior to onset of the hold phase. In the remaining 2 neurons, the response to the ramp stimulus was ambiguous; in one case, the GCFR was noisy with no apparent trend and in another, the firing rate decreased and increased within the duration of ventrad ramp movement.

For instance, in the neuron shown in **Figure 2A**, the firing rate increased during ventrad ramp movement (bluegreen trace) and decreased during dorsad ramp movement (purple trace) (**Fig 2A i, bottom panel**). Thus, this neuron showed directional sensitivity. Moreover, the neuron responded with higher firing rates for dorsad hold position with slight adaptation and lower firing rates for ventrad hold positions (also see **Supplementary Figure 1B**). During the hold phase, the mean steady firing rate varied linearly as the angular position at which the antenna was held (p ≤ 0.01, r^2^ ≥ 0.8, Wilcoxon rank sum test, **Fig 2 B**), demonstrating their sensitivity to angular position (also **Supplementary Figure 1**).

In 4 out of 20 recordings (N=18 moths), the firing rate of JO neurons varied linearly with angular position of the flagellum (**Fig 2C**). Of these, two neurons increased their firing rate, and two decreased their firing rate in response to varying dorsad hold positions of the flagellum. In our experiments, neurons were not labelled, and hence we could not determine their position within the Johnston’s organ relative to the stimulus, or if dorsad or ventrad ramp movements corresponded to stretch or compression of the JO mechanoreceptors. In the remaining 16 neurons, the firing rates during the hold phase were not significantly different (p>0.05) from each other or else they dropped to baseline levels after the initial rise during the ramp phase, and hence did not show position-sensitivity.

### JO neurons encoding angular velocity of the flagellum

We next examined if JO neurons were sensitive to angular velocity of the flagellum by subjecting them to a protocol in which the flagellum was moved to the same angular position, *albeit* at variable angular velocities broadly ranging from as low as ∼0.1°/s to as high as ∼4°/s.

Figure 3A shows recordings in response to the above protocol for the same neuron as Fig 2. Here, the antenna was subjected to 7 different angular velocities of which 4 representative stimuli and their responses are shown (**Fig 3A**). In these 4 stimuli, the angular displacement was ∼0.7° but the angular velocities were 0.46, 0.86, 1.68, 3.16 °/second respectively (**Fig 3A**, **top and middle panels**). We estimated their angular velocity by fitting a line through the ramps at the beginning of the stimulus (**Fig 3A ii**, top panel, see methods for the angular velocity estimation). This neuron fired at a maximum rate (∼145 Hz) for the maximum angular velocity presented (∼3.16 °/s, purple trace), and at proportionally lower rates for the lower angular velocities (**Fig 3A i**, **bottom panel**; magnified in **Fig 3A ii**, bottom panel).

**Figure 3.**
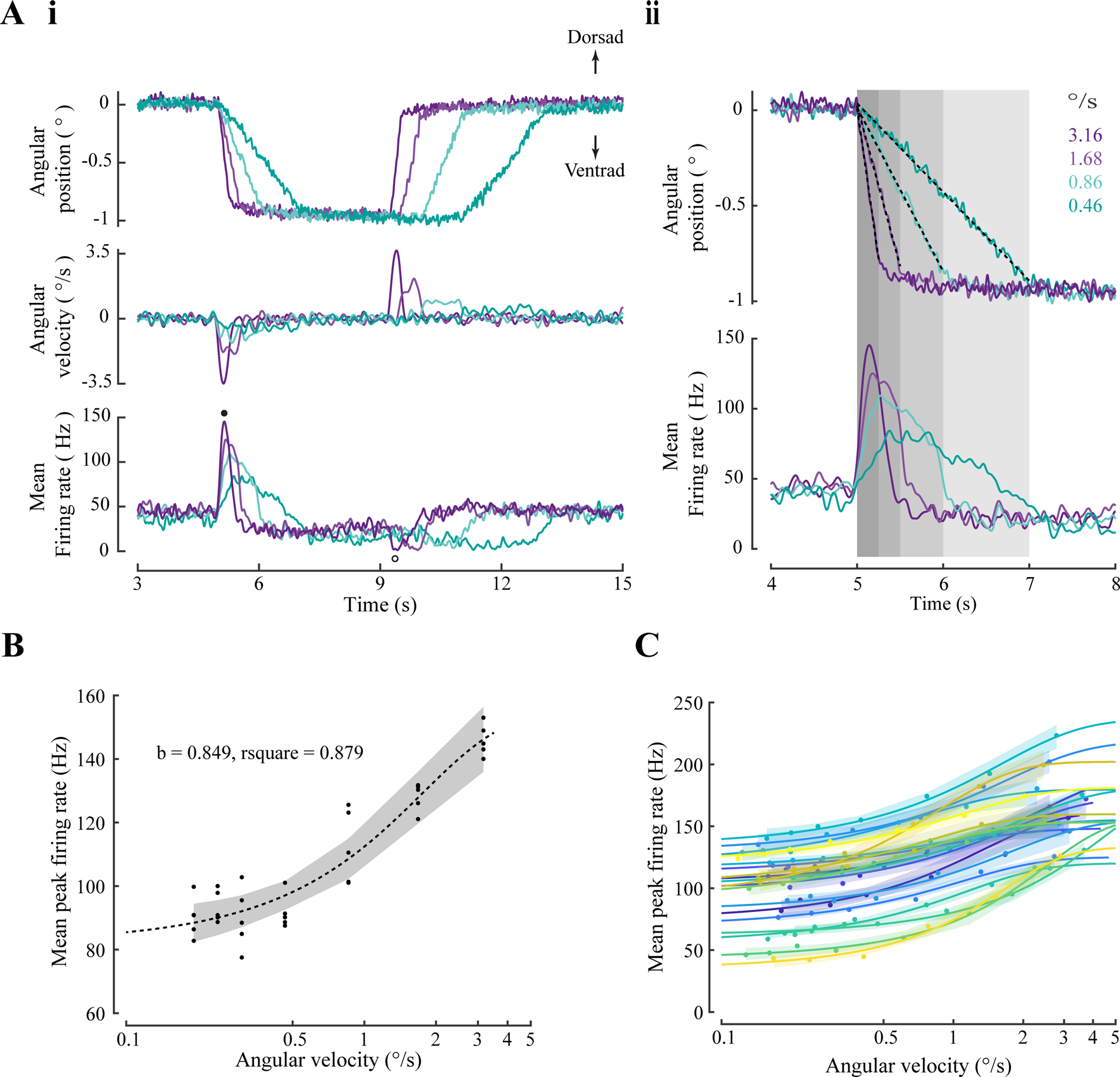
Velocity-encoding neurons. (**A i**) (*top*) Mean angular position stimulus delivered to flagellum (5 trials per protocol). The velocity stimulus varied from 0.2 °/s to 3.1 °/s. (4 out of 5 protocols shown for clarity; bluegreen to purple traces with velocities: 0.46, 0.86, 1.68, 3.16 °/second); (*middle*) Angular velocity profile; (*bottom*) Average firing rate obtained from 5 trials for each velocity stimulus. Increase and decrease in firing rate corresponding to ventrad and dorsad ramp movement are marked by closed and open circles respectively. (**A ii**) A detailed view of the peak firing rate during the constant angular velocity stimulus. The grey boxes mark the duration of the ramp stimulus. The dotted black line is the linear fit performed on the ramp to estimate angular velocity. (**B**) Mean peak firing rate response to angular velocity is modelled by a logistic function. *Dashed line*: logistic fit, *shaded region*: 95% confidence interval. The x-axis is on log scale. (**C**) In 19 out of 27 neurons, r^2^ ≥ 0.8. The solid lines are the logistic fits for individual neuron and shaded region shows the 95% confidence interval of the fit.

Thus, these neurons were sensitive to the angular velocity of the flagellum. We employed 4 different models to ascertain the relationship between the peak firing rate and angular velocity (details in the **Supplementary material**). Based on this comparison, we chose a logistic fit as the most appropriate to explain the relation for 19 out of 27 neurons (**Fig 3C**, r^2^ ≥ 0.8), including the neuron in Fig 3A (**Fig 3B**, r^2^ = 0.88).

Ventrad (downward) ramp movement of the flagellum increased the firing rate (closed circle), whereas dorsad (upward) ramp movement decreased the firing rate (open circle), thus demonstrating directional sensitivity of the JO.

### JO neurons exhibiting adaptations to angular position and velocity

When the neuronal activity of a sensor decreases with time for a constant stimulus, the phenomenon is termed ‘adaptation’ (Adrian and Zotterman, 1926; Thorson and Biederman-Thorson, 1974; for a more recent review see Benda, 2021). Typically, this decrease is exponential, and may be characterized using the time constants of these exponential decays (see methods).

JO neurons exhibited diverse adaptation dynamics during the ramp-and-hold protocols. Figure 4A shows the responses of 7 different neurons to similar stimuli (**Fig 4A**). The inherent adaptation of the responses to these stimuli varied widely. These responses are baseline subtracted for the ease of visual comparison. For instance, the two responses highlighted in **Fig 4A, bottom panel** (blue and orange traces) show different adaptation dynamics to ramp movement and hold position. The blue trace shows a slow adaptation to constant angular velocity compared to the orange trace, captured by adaptation time constant of 11.2 ms versus 1.6 ms (**Fig 4B ii, highlighted in pink**). During the hold position stimulus, the blue trace adapts faster than the orange trace, captured by adaptation time constant of 9.6 versus 37.3 ms.

**Figure 4.**
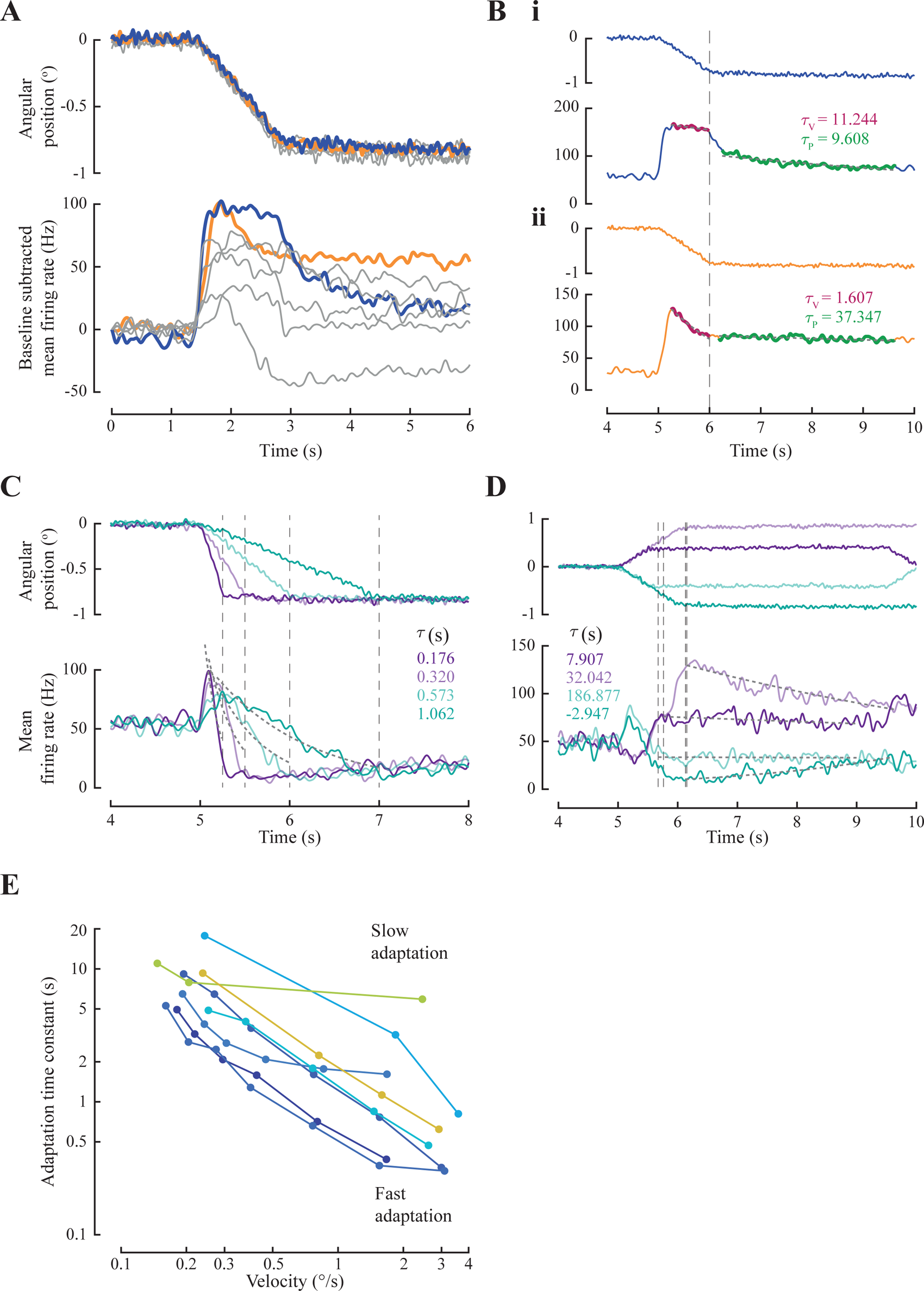
Neurons exhibit adaptation to angular velocity and position. (**A**) Response of 7 neurons to similar stimuli with steady-state positions of -0.84° ± 0.03° (mean ± SD), showing a wide range of adaptation dynamics exhibited by JO neurons. Response of 2 neurons are highlighted in blue and orange for comparison. The baseline firing rate is subtracted for the ease of visual comparison of the nature of adaptation. Here, the ramp stimulus starts at 1.5 s. (**B**) Adaptation is quantified as the time constant of the exponential decay of the firing rate. The response of the two neurons highlighted in A, retaining the colour code. The dotted line marks the end of the ramp stimulus and start of hold stimulus. (**B i**) (*top*) Ramp-and-hold stimulus. (*bottom*) Response of a neuron that shows slower adaptation to angular velocity but faster adaptation to constant position. (**B ii**) (*top*) Ramp-and-hold stimulus. (*bottom*) Response of a neuron that shows fast adaptation to velocity and slow/no adaptation to constant position stimulus. (**C**) Exponential fits and adaptation time constants to responses for 4 different angular velocities for the neuron in Supplementary figure 2A. Fits are shown with grey dotted lines. (r^2^ > 0.75) (**D**) Exponential fits and adaptation time constants to responses for 4 different angular positions for the same neuron in C. Fits are shown with grey dotted lines. (r^2^ > 0.5) (**E**) Responses of 8 neurons (on a log-log scale) show a decrease in adaptation time constant with an increase in angular velocity. 8 out of 26 neurons remained after eliminating the trials that had a fit with r^2^ < 0.75 and neurons that had less than 3 quantifications of time constant or positive time constants.

Thus, individual JO neurons had different adaptation properties that determined their response dynamics. We quantified the adaptation in response to the two parts of the ramp-hold stimulus with two separate single exponential fits (see methods) (**Fig 4B**). τ corresponds to the adaptation time constant. Note that a lower value of time constant indicates faster adaptation, and a larger time constant indicates slower adaptation. The subpanels **Fig 4B (i-ii)** present data from 2 different neurons that adapted to angular velocities (ramp phase) and positions (hold phase) to different extents.

Figure 4C shows the adaptation of firing rate for the neuron that linearly encodes angular positions (**Supplementary Fig 1**), during the ramp stimuli for constant hold position and varying angular velocities. Figure 4D shows the adaptation during varying hold positions, while the ramp velocity was held constant. The values of adaptation constants in these protocols show that even within a single neuron, the adaptation dynamics varied with the value of angular velocity and angular position of the flagellum, i.e. adaptation was faster for larger values of angular velocity and position. As the angular velocity reduced, the firing rate took longer to adapt to the baseline firing rate, as reflected in greater time constants (**Fig 4C**, **bottom panel**). For larger angular deflections, the adaptation time constants were smaller, recapitulating faster adaptation. Also, during the hold stimuli, the adaptation was slower for positions closer to the baseline (**Fig 4D**). Across 8 neurons, the time constant consistently reduced with an increase in angular velocity (**Fig 4E**).

### JO neurons that are direction-sensitive and history-dependent

A prominent feature in most neurons was their sensitivity to the direction of ramp movement. Because the movement stimulus in the experiments reported here was uniaxial, the directional responses recorded here were restricted to the dorso-ventral movement of the flagellum. When stimulated with the stair protocol, 55 of 59 recorded neurons exhibited increased activity exclusively to either dorsad or ventrad ramp flagellar movement, indicating a strong directional preference. Opposite to the preferred direction, the firing rate decreased. Such response reversal was observed whenever stimulus changed direction (e.g. **Fig 2A, 3A, 5Ai** and **Bi, Supplementary Fig 1A,C**). In the remaining 4 neurons, the response was ambiguous due to variability in the response. Moreover, this directionality was exhibited for both ramp movement and hold position in opposite directions. For example, in Figure 2 and Supplementary figure 1A, neurons showed preference to ventrad ramp movement, but dorsad hold positions. Because these recordings were from primary mechanosensory neurons with low latency responses, the reduced response likely reflects a reduction in spiking activity (membrane potential and raster plots in **Fig 5A i and 5B i**) rather than active inhibition.

**Figure 5.**
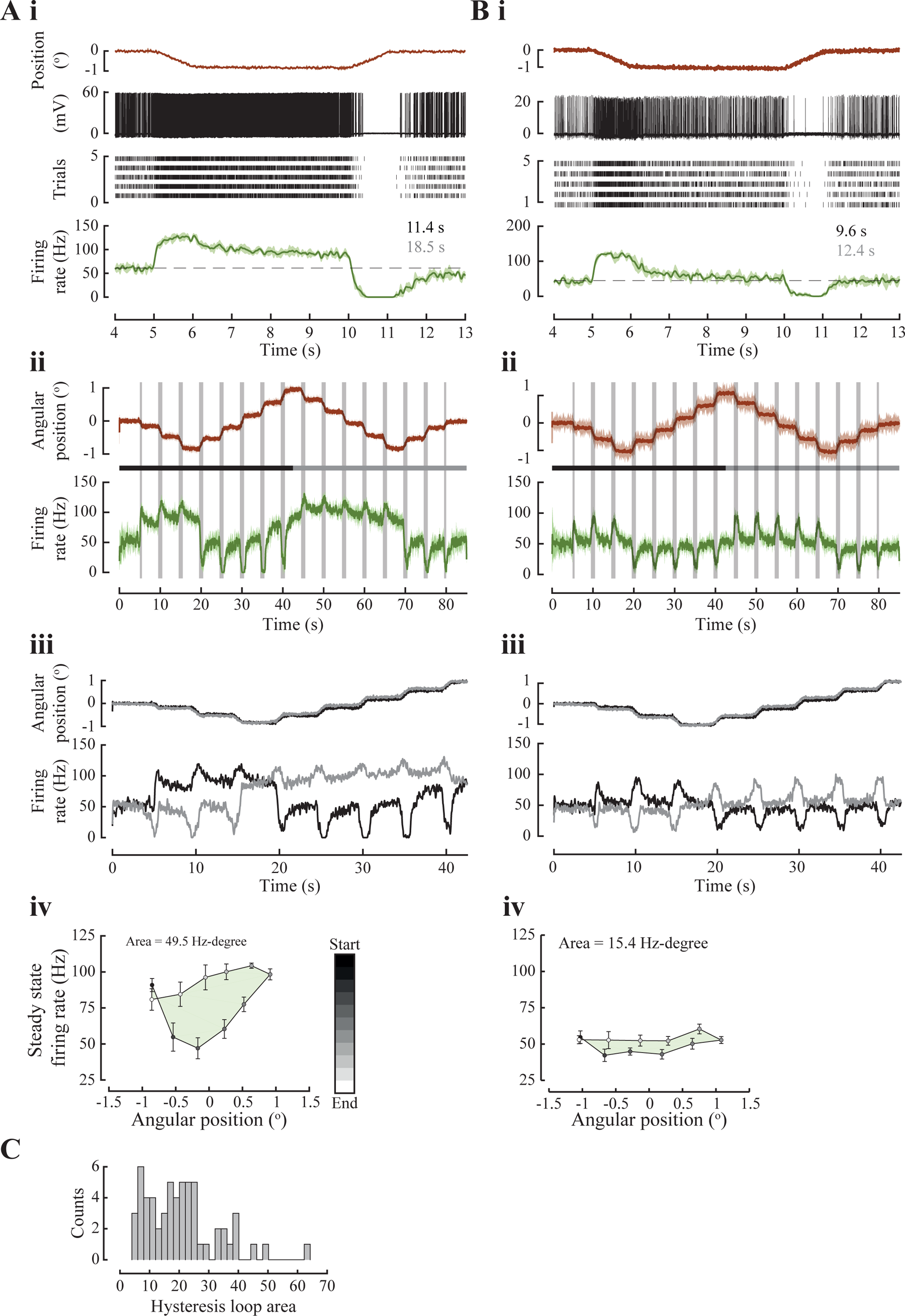
Direction-sensitive responses of 2 neurons with low and high hysteresis. (A, B) (**i**) (*top-bottom*) Ramp-and-hold stimulus; Membrane potential of the first trial showing absence of spikes during dorsad ramp movement of the flagellum; raster plot showing response in 5 trials. Note the increased spike density in response to ventrad ramp movement and the stark absence of spikes during dorsad ramp movement; the response shows a peak firing rate for ventrad ramp movement, an adapting firing rate during the hold phase and reaches 0 firing rate during dorsad ramp movement. Adaptation constants for adaptation of response to constant angular velocity (in black) and constant angular position (in grey). (**ii**) (*top*) Stair stimulus protocol (see methods). (*bottom*) Firing rate across trials (mean ± SD). Grey bars mark the ramps between hold positions. (**iii**) Same as (**ii**) with the stimulus and response cut midway and overlayed to compare the responses to instances of the same positions within the protocol. In black, the stimulus and response from the beginning of the stimulus protocol to the midway point (shown with a black line below the stimulus trace in (**ii**). In grey, the stimulus and response from the midway point to the end of the stimulus protocol (shown with a grey line below the stimulus trace). (**iv**) Hysteresis in mean steady-state firing rate (mean ± SD) response to similar angular positions. The colour of the circle marks the sequence of positions followed in the stimulus protocol. Initial and final 3 hold positions are not considered in the quantification of the hysteresis. The value in the top left is the area of the hysteresis loop. (**C**) Histogram of area of the hysteresis loop from 59 neurons, showing that many JO neuron exhibit hysteresis.

To observe how direction-sensitivity influenced the response to constant position stimuli, we designed a *stair protocol* in which the antenna was held at various positions while keeping constant the step size and the angular velocity during the transition. Here, the same hold positions were attained during dorsad and ventrad excursions. We recorded response of 59 neurons from 32 moths to the stair protocol. For instance, **Figs 5 A ii and 5 B ii** show the response of 2 neurons to the stair protocol. In both neurons, there is a brief increase in activity during every ventrad ramp movement, and a brief decrease during every dorsad ramp movement. The response adapted to the steady-state firing rate when hold position was constant (**Fig 5Ai, ii and 5Bi, ii**).

Reversing and replotting the responses to upstairs and downstairs stimuli with overlapping hold positions (**Fig 5A iii, 5B iii**), shows that the steady-state firing rate encoding the hold position is significantly different for rising *versus* falling stimuli for the same absolute position for the neuron in Fig 5A. In contrast, the response to hold-positions for neuron in Fig 5 B consistently returned to the baseline. For both neurons, the preceding movement history influences response to hold positions. The neuron in Fig 5A exhibits a strong asymmetry in responses to ventrad and dorsad ramp movements.

History-dependence, which causes hysteresis, may be quantified as the area of the loop formed by the firing rate response to one cycle of positions starting from the most ventrad position to the most dorsad position and back. Thus, the neuron in Fig 5A exhibits a large hysteresis in the steady state firing response with an area of 49.5 Hz-degree (**Fig 5A iv**). In contrast, the neuron in Fig 5B has lesser hysteresis, with area of 15.4 Hz-degree (**Fig 5B iv**).

The second source of hysteresis is the adaptation time constant. The first neuron (**Fig 5A i**) adapted more slowly to angular velocity (τ_v_ = 11.4 s) and to angular position (τ_p_ = 18.5 s) (**Fig 5A i**), while the second neuron exhibited a slightly faster adaptation to both velocity and position (τ_v_ = 9.6 s, τ_p_ = 12.4 s) (**Fig 5B i**). Across 59 neurons we see a wide range of hysteresis values ranging from 4.67 to 64.11 Hz-deg (**Fig 5C**).

### JO neuron showing position-dependent response to angular velocity

In 20 neurons (N=18 moths), we provided ramp-and-hold protocols which recorded responses to both variable ramps and variable hold positions. Of these, 1 neuron responded strongly to ventrad ramp movements, but remained largely inactive to dorsad ramp movements of the flagellum (Fig 6 **A, B**). Its peak firing rate in response to varying ramp velocity (but constant hold position) varied in a sigmoidal manner (**Fig 6C**). In this neuron, the peak firing rates were greater when the flagellum moved from a more dorsad to zero position (**Fig 6B**). The responses to ventrad ramp movement from zero position was significantly lower than the responses to ramp movement in the same direction but from more dorsad hold positions (**Fig 6D**). Thus, for this neuron, the peak firing rates in response to ramp movements depended strongly on the value of its hold position. In contrast to this neuron, in a majority of other 19 recordings, the peak firing rate for ventrad ramp movement did not depend on their hold position.

**Table 1:** Summary of responses of all position-, velocity-, and direction-sensitive neurons. Responses of 9 neurons were excluded as they did not significantly change their activity in response to velocities, positions, or start/end of deflections.

| Response type | No. of neurons<br>(No. of moths) |
| --- | --- |
| Ventrad velocity sensitive | 19/27 (26) |
| Ventrad hold position and ventrad velocity sensitive | 2/20 (18) |
| Dorsad hold positions sensitive and ventrad velocity | 1/20 (18) |
| Sensitivity to dorsad hold positions and the start of ventrad ramp movement | 1/30 (26) |
| Sensitivity to onset and offset of movement in either direction | 1/30 (26) |
| Ventrad ramp movement | 56/60 (32) |
| Dorsad ramp movement | 1/60 (32) |
| Dorsad hold positions | 3/20 (18) |
| Ventrad hold positions | 2/20 (18) |

**Figure 6.**
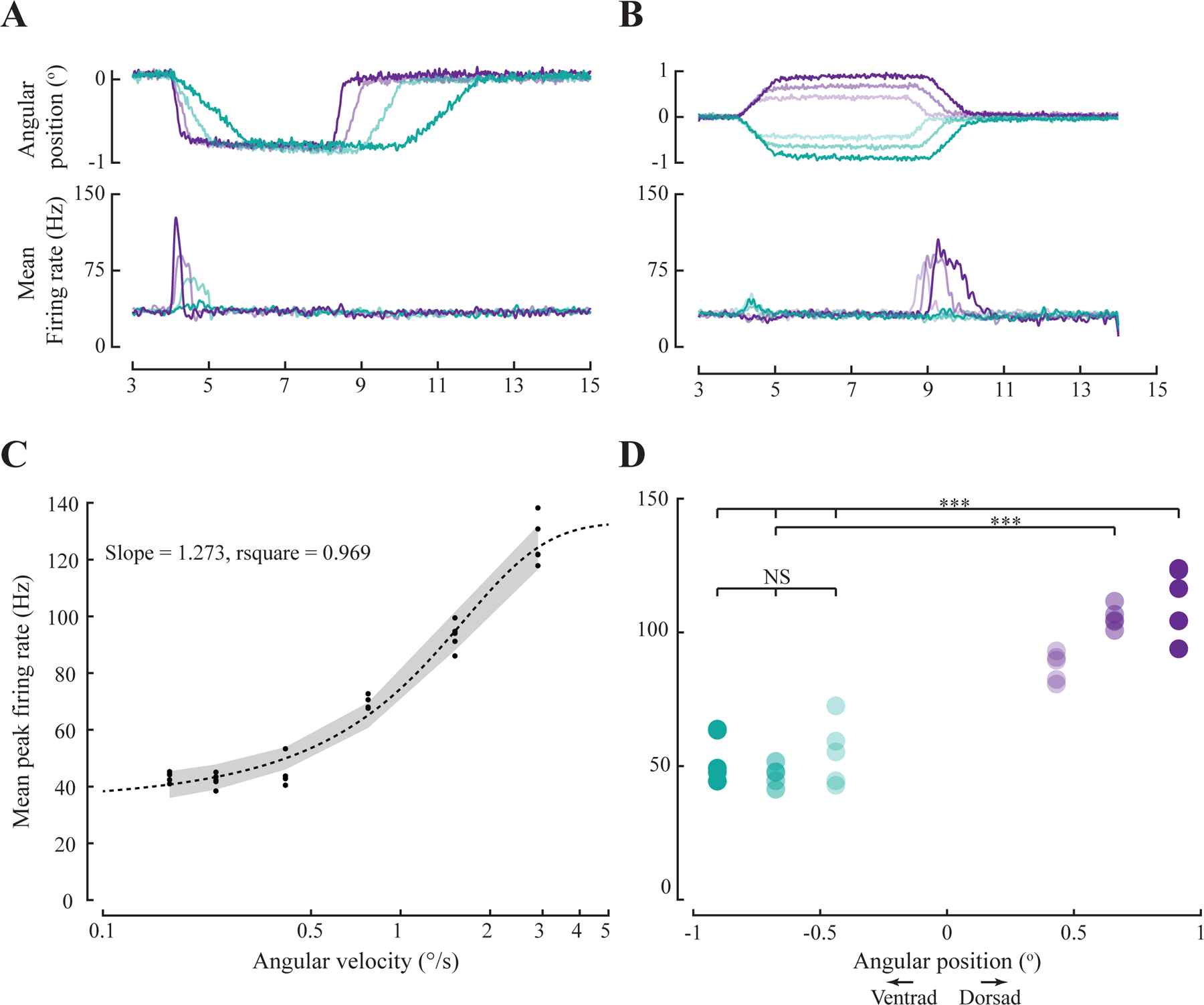
Position-dependent response to velocity. (A) (*top*) Ramp-and-hold stimulus with variable ramp velocities (0.2 to 2.8 °/s); (*bottom*) Corresponding responses in terms of mean firing rate (color-coded). (B) (*top*) Ramp-and-hold stimulus with variable hold positions (-0.9 to 0.9 °/s); (*bottom*) Corresponding responses in terms of mean firing rate (colour-coded). (C) Logistic fit of peak firing rate versus angular velocity (r^2^ = 0.89), suggesting that the neuron encodes velocity (D) Comparison of peak firing rate in response to ventrad ramp movement from positions 0.4, 0.66, and 0.9 ° at the same angular velocity of ca. 0.76 °/s.

## DISCUSSION

The experiments described in this study test the hypothesis that mechanosensory Johnston’s organs encode the angular position and angular velocity, which are the initial two components in the series decomposition of any time-varying mechanical stimulus. We performed intracellular recordings from axons of scolopidial units of the JO at the base of the antenna, while selectively varying the angular position and angular velocity of the flagellum using ramp-and-hold stimuli. Angular position was altered by moving the antennae to different hold positions, whereas angular velocity was varied by moving the antennae along ramp motions of different slopes. We recorded from 60 neurons, from 32 moths. We also detected the actual movement of the antenna using a Hall effect sensor placed very close to the location of the JO neurons.

Most neurons exhibited some baseline activity at the outset, likely due to residual stimulus at resting position. In such cases, the response to a delivered stimulus was evident as a change relative to this baseline activity. Also, most neurons increased activity during a ventrad ramp movement of the antenna, suggesting a strong response in the direction of gravity. In all neurons, the firing rate changed in response to antennal movement. These recordings show that JO neurons encode angular position and angular velocity, in addition to other properties such as directional encoding, and rapid or slow adaptations to either angular velocity or position stimuli or both.

### JO responds to ventrad ramp movement of the flagellum, in the direction of gravity

A vast majority (56 out of 60) of the neurons responded to ventrad ramp movement of the antenna (**Table 1**), whereas their activity was inhibited by dorsad ramp movement. In the resting state and also when the antenna is positioned prior to onset of flight, the JO primarily experiences the weight of the flagellum in the direction of gravity, which suggests the intriguing possibility that the antenna serves to sense the direction of gravity. In many insects including Drosophila etc, the antenna has previously been hypothesized as being a gravity sensor (Kamikouchi *et al*., 2009), but other data contradict this possibility (Kladt and Reiser, 2023).

The observation that JO responds strongly to ventrad ramp movement of the antenna is consistent with the hypothesis that JO serves to detect gravity. Because the preparation described here was severely restricted to enable recordings from axons of single neurons, it was not possible within the scope of these experiments to test this function.

### JO neurons encode angular position and angular velocity

Our study showcases individual scolopidial units of the JO that encode both angular position and/or angular velocity. As the first two components in the series expansion of any arbitrary, time-varying stimulus, the encoding of position and velocity ensures that the JO can capture a major component of the antennal motion ranging from various low frequency stimuli such as airflow or gravity (e.g. Gewecke 1974, Khurana and Sane 2016, Natesan *et al*., 2019, Kamikouchi *et al*. 2009, Yorouzu *et al*. 2009, Suver *et al*. 2019), as also high frequency stimuli such as antennal vibrations at wing beat frequency (Sane *et al*., 2007; Dieudonne, Daniel and Sane, 2014). When insects including moths, honeybees, bumblebees etc. are deprived of these stimuli through flagellar ablations, their flight control is compromised. The Johnston’s organ also plays a significant role in antennal positioning behaviour during flight in tethered hawkmoths (Natesan *et al*., 2019) and freely-flying honeybees (Khurana and Sane, 2016). Under constant airflow conditions, elimination/reduction of inputs to the JO by gluing of the pedicel-flagellum joint eliminates the modulation of antennal positioning in both hawkmoths and bees. Angular position and velocity encoding in combination with the direction sensitivity of the JO neurons may be essential for antennal positioning. The inputs from JO are shown to be critical for head stabilisation, which in turn is essential for stable flight (Chatterjee *et al*., 2022). The ability of some neurons to encode both angular position and velocity (Fig 2**, 3**) also points to their functional versatility.

### JO neurons are range-fractionated

Sensory neurons must contend with the trade-off between range and sensitivity. In many sensory organs including diverse eyes (Land and Nilsson, 2012), hearing systems (Göpfert and Hennig, 2016), proprioceptive organs (Barth, 1964) etc, this trade-off is resolved by distributing range over multiple units that compose a sensory organ, while ensuring that each of those units is narrowly tuned to a specific frequency range. Previous research has shown range fractionation in frequency encoding by JO neurons in hawkmoths (Sane *et al*., 2007; Dieudonne, Daniel and Sane, 2014) and, more recently, in flies (Patella and Wilson, 2018). Similarly, the mechanosensory neurons that constitute the Femoral Chordotonal organs in insect legs also exhibit range fractionation of leg angles over which individual neurons are sensitive to tibial movements (Matheson, 1992). The mechanosensory cercal hairs in crickets are range fractionated by the length of the sensilla (Shimozawa and Kanou, 1984). Many studies in the field relating to mechanosensors leave open the possibility that the mechanical properties, geometry or location of the sensory units within an organ may play a key role in range fractionation, in addition to selective neuronal tuning properties.

In our experiments, the peak firing rate of JO neurons encoded angular velocity were modelled as a sigmoidal (logistical) function (**Fig 3C**). Because the stimulus range in these experiments was only about 20 fold (∼0.2 to 4 °/s), we were unable to explore the possibility of power law relationships between the mechanical deflection of the antennae and the corresponding JO neuron responses, which typically require a stimulus range of at least two orders of magnitude. The practical difficulties in achieving such a range include, on the one hand, the ability to deliver very fine and precise mechanical stimuli, while on the other to deliver stimuli of a large magnitude. Such power law relationships have been previously suggested as being important in range compression by sensory neurons to enable single neurons to encode over a large range of stimuli (e.g. Thorson and Biederman-Thorson, 1974).

The various JO neurons characterized in this study vary in their sensitivity to delivered range of velocities, represented by varied range of slopes (**Fig 3C**) which determines the range of velocities over which they encode. Consistent with previous data on *Manduca sexta* (Sane et al, Dieudonne et al), JO neurons in *D. nerii* also show range fractionation, which enables the antennae to serve a multimodal role ranging from sensing steady or low frequency signals as required for airflow detection and gravity sensing (Kamikouchi *et al*., 2009) to sensing high-frequency signals as required for vestibular feedback (Sane *et al*., 2007).

### JO neurons display adaptation of response

Adaptation is a common feature across sensory neurons, interneurons, and motor neurons. Here too, a power law (see methods) with fractional exponent has been proposed to capture adaptation, which may result from distributed relaxation processes (Thorson and Biederman-Thorson, 1974), distributed viscoelastic coupling (Chapman, Mosinger and Duckrow, 1979), and viscoelasticity of microtubules (Kuster, French and Sanders, 1983). Power law adaptation has been shown at the stage of coupling between stimulus and the receptor (Loewenstein and Mendelson, 1965; Chapman, Mosinger and Duckrow, 1979), during transduction (Nakajima and Onodera, 1969a, 1969b) and action potential generation (e.g. in femoral tactile spine in cockroaches, French, 1984; French, 1984b). In chordotonal organs, adaptation may occur at the transduction or spike generation stage (Field and Matheson, 1998).

Because we observed distinct adaptations to angular velocity and angular position stimuli, these two adaptations were separately characterized, using two exponents corresponding to the different time constants for velocity and position, respectively. The adaptation time constant was smaller for higher velocities of the stimuli and larger for lower velocities. Since peak firing rate in response to angular velocity increases with angular velocity, the adaptation time constant may be related to the peak firing rate (**Supplementary figure 3**). Firing rate may adapt faster following a higher firing rate due to faster depletion of active channels involved in spike generation.

In sensory systems, adaptation performs a high-pass filtering of stimulus (Benda, 2021), thereby increasing the frequency threshold for activation. This process ensures sensitivity to small fluctuations riding over a sustained low-frequency stimulus. When combined with other non-linear phenomena, adaptation may help in ensuring stimulus selectivity and reduction of noise. For instance, adaptation to mean has been described in the fly auditory system where the JO neurons correct for the background noise, thus remaining sensitive to courtship song (Clemens, Ozeri-Engelhard and Murthy, 2018). In hawkmoths, we propose the hypothesis that adaptation helps keep the JO sensitive to minute perturbations when flying in a constant airflow condition (Natesan *et al*., 2019), or performing an aerial turns (Sane *et al*., 2007; Dahake *et al*., 2018).

### Hysteresis in encoding position

Besides the encoding features introduced by adaptation, it is evident from the responses to stimuli in the stair protocol, adaptation along with a directional response to stimuli brings in a strong element of nonlinearity, including hysteresis, into sensory encoding. Hysteresis in a neural response is the dependence of neural activity on the history of stimulus, in addition to the current stimulus (Hatsopoulos, Burrows and Laurent, 1995). In the data presented here, the observed difference in steady-state firing rate in response to the same positions depended on the stimulus history, resulting from both adaptation and the directional response property of the JO neurons. Hysteresis behaviour has been reported in locust Femoral Chordotonal organs (Burns, 1974; Zill, 1985; Matheson, 1992), trochanteral Campaniform sensilla in stick insects (Hofmann and Bӓssler, 1986) and in cockroach tibial campaniform sensilla (Ridgel *et al*., 2000). Such hysteresis may occur due to asymmetries in the morphology of the sensor, biomechanical properties of the interstitial tissue, neural adaptation, or a combination of these factors. Such history-dependence is surprising because it means that the primary mechanosensory neurons provide feedback not only about the instantaneous stimulus state but also about how that state was attained. The implications of such history dependence for sensing of natural stimuli remain largely unexplored.

### The similarity of responses to FeCO and Campaniform sensilla

The response properties of JO neurons are analogous to those observed in FeCO in insect legs. Similar combinations of position, velocity, acceleration, and direction encoding have been demonstrated in FeCO of stick insects and locusts (Hofmann and Koch, 1985; Hofmann, Koch and Bässler, 1985; Zill, 1985; Buschges, 1994). The stretch receptor organ in caterpillars of a hawkmoth also encodes position and velocity (Simon and Trimmer, 2009).

Despite significant differences in structure and spatial distribution on the body, force encoding in campaniform sensilla in cockroaches (Ridgel *et al*., 2000; Zill, Büschges and Schmitz, 2011) and encoding of leg bending in stick insects (Hofmann and Bӓssler, 1986) exhibit similar responses in terms of encoding angular velocity of movement or rate of change of force, encoding constant position or constant force, adaptation, and hysteresis. These response characteristics are comparable to responses in coxo-basal chordotonal organs in the legs of crabs (Bush, 1965) and intraspinal mechanosensory neurons in lamprey (Massarelli *et al*., 2017).

The similarities across various mechanosensory organs, despite their structural and functional diversity, suggest that the properties of the mechanosensory neurons are largely similar. However, specific encoding by these sensors may be determined by their location, biomechanics (Sane and McHenry, 2009; Barth, 2019), and associated mechanoreceptors.

The stimulus experienced by the sensory neuron is filtered by the physical structure of the mechanosensor (Sane and McHenry, 2009; Barth, 2019). A mechanical constraint on the movement causes a directional response in sensors, as seen in cercal filiform hairs in crickets and trichobothria in spiders. The orientation, relative to the movement axis of tibial campaniform sensilla in cockroaches (Zill, Moran and Varela, 1981), wing campaniform sensilla in flies (Dickinson, 1992) and slit sensilla in spiders determines the strain patterns that are encoded (Barth, 2019). The mechanical properties determine the frequency that reaches the sensory organ, for example, cercal filiform hairs in crickets, lateral line system in fish, and tympanal organs in various insects (Sane and McHenry, 2009). A comparative study of the biomechanics of these mechanosensors, along with recordings of receptor potentials will help us identify the key elements that sculpt the unique functionalities of these mechanosensors.

## Conclusions

This study provides evidence for encoding of angular position, angular velocity and direction of flagellar movement by Johnston’s organ in hawkmoths. The JO neurons exhibit adaptation to either/or angular positions and velocities and history dependent response to angular positions. A vast majority of the neurons increase their activity in response to ventrad movements but their activity is suppressed for dorsad movements, which suggests direction sensitivity in the direction of gravity. The similarities in encoding properties to other mechanosensors across arthropods highlight the importance of biomechanics of these sensors that allow encoding of diverse mechanosensory stimuli.

---

**Supplementary figure 1.**
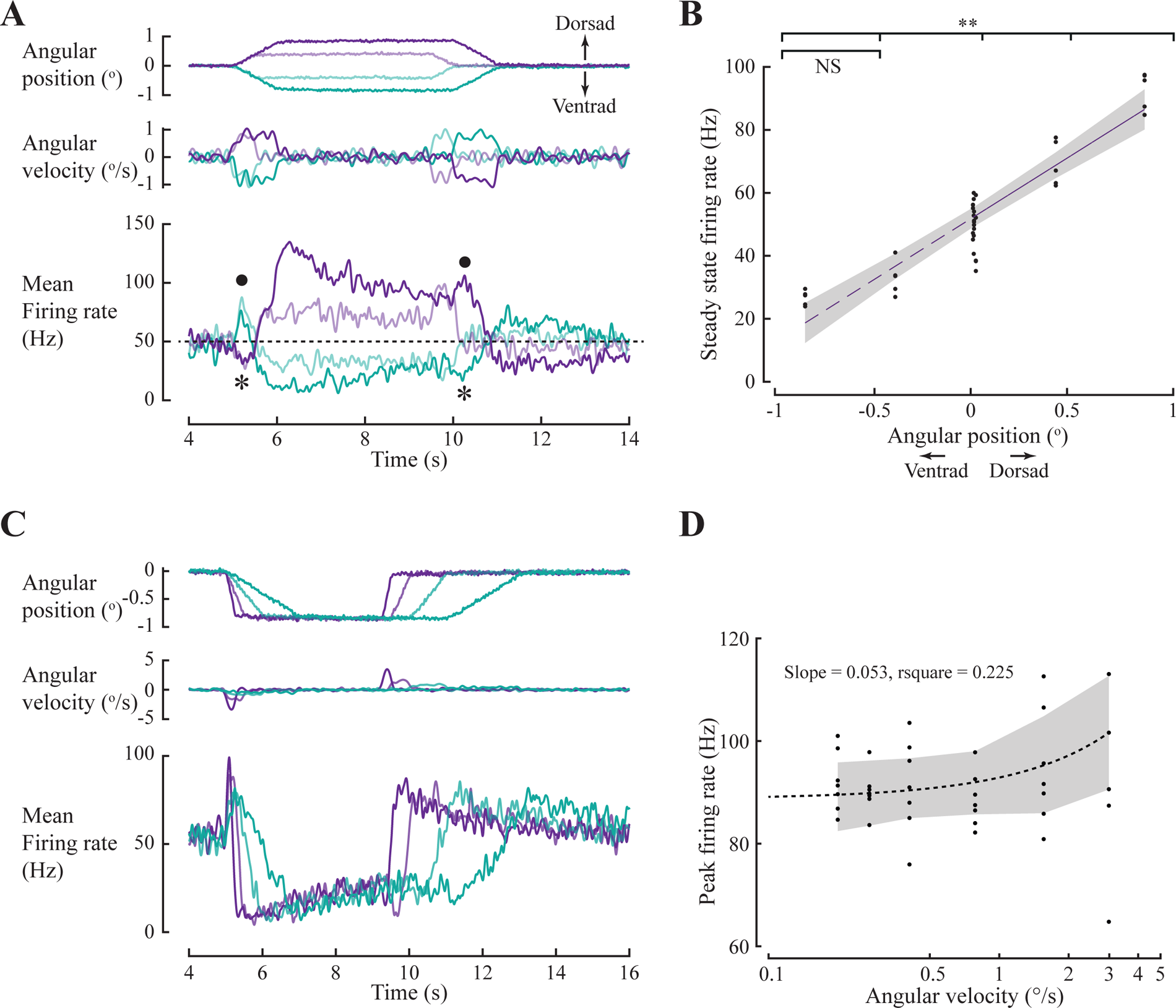
Ventrad ramp-movement and position-sensitive neuron. (**A**) Same as Fig 2A. This neuron exhibits a brief increase in firing rate when a ventrad ramp movement is initiated (dot) and a brief dip in firing rate when movement is initiated in the dorsad direction (star). After this transient response, the response dynamics change within the ramp duration. (**B**) The steady-state firing rate linearly increases with the angular position of the flagellum (r^2^ > 0.8). Stars indicate p-value < 0.01 (Wilcoxon rank sum test). (**C**) Same as Fig 3A. (**D**) peak firing rate does not significantly vary at different velocities of movement. (r^2^ = 0.24).

**Supplementary figure 2.**
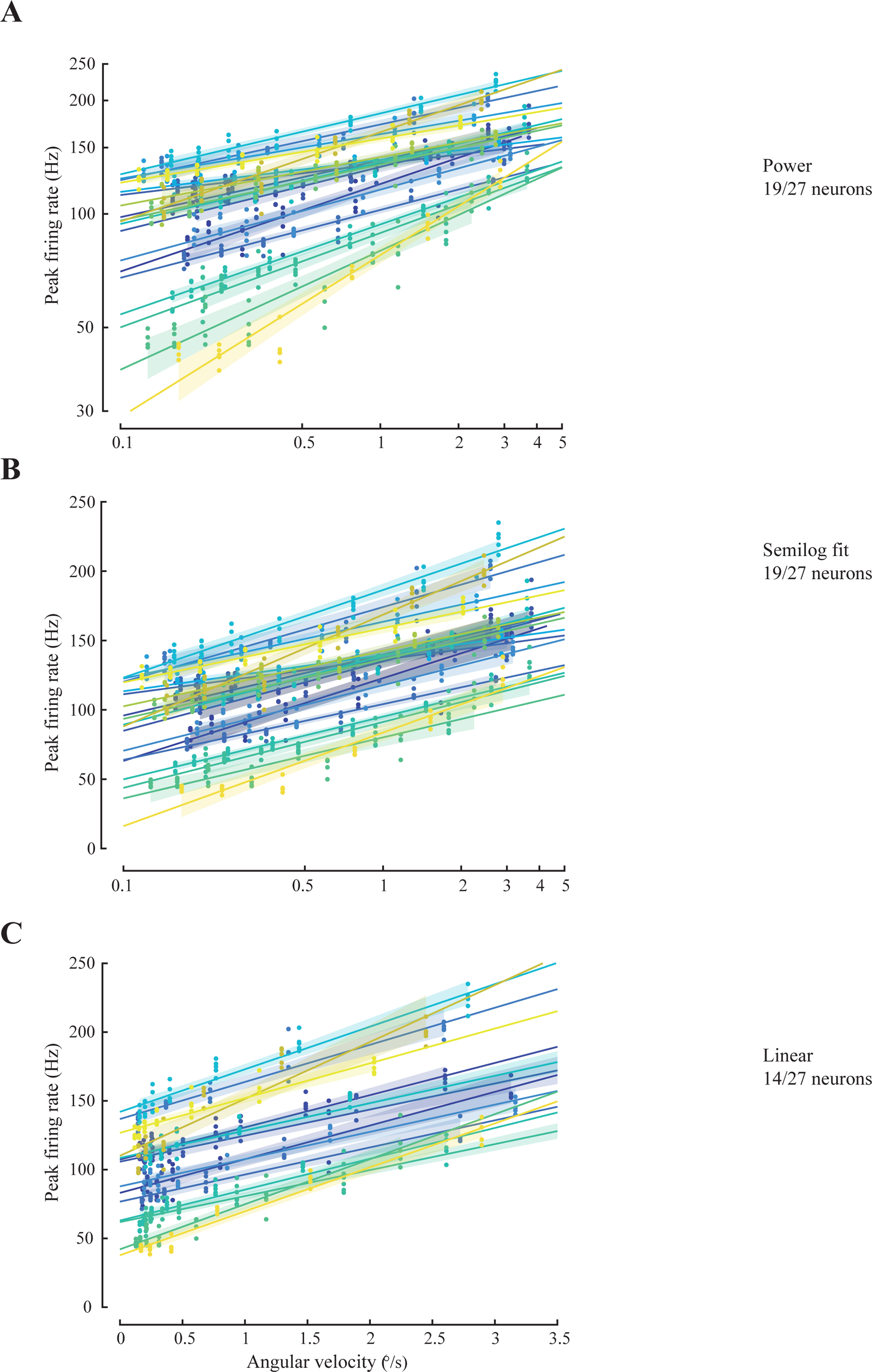
Peak firing rate response versus angular velocity fit to other models. Response of 19 of 27 neurons explained (A) Power law; (B) Semi-log fit; (C) Linear fit. (r^2^ ≥ 0.8 for all). Shaded region around the fit is 95% confidence interval around the fit.

**Supplementary figure 3.**
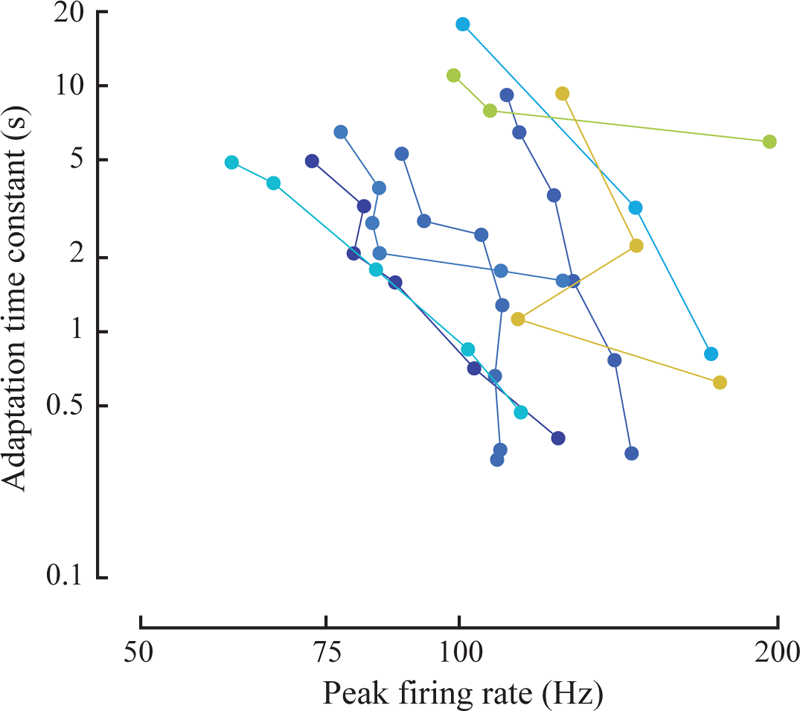
Adaptation time constant versus peak firing rate elicited by different angular velocities. Same neurons as Fig 4E.

## Supplementary Information

To capture the relationship between peak firing rate and the angular velocity we fit multiple models suggested in the literature and compare how much data are explained by each model. 3 out of 30 neurons where the response to ramp stimulus was not clear, were excluded from this analysis.

*Linear fit.* To test the first order relation, we fit a first degree polynomial.

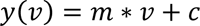

where,

*y*(*v*) = Peak firing rate, *v* = Angular velocity of flagellar movement,

*c* = Y-intercept.*Power law fit.* A power law has been used to model the relation between flexion and extension velocity of the femoral chordotonal organ (Matheson, 1992) and the relation between the rate of change of forces on the tibial campaniform sensilla (Ridgel *et al*., 2000; Zill, Büschges and Schmitz, 2011). Here we have used it to show the relation between peak firing rate and the angular velocity of movement. The power law is given by:

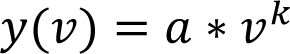

which can be written as,

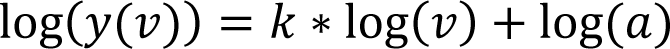

where,

*y*(*v*) = Peak firing rate, *v* = Angular velocity of flagellar movement,

log(*a*) = Intercept, and

*k* = Slope of the line on a log-log plot.

*Semilog fit.* The range of angular velocities and firing rates were speculated to be very narrow for using a power law. So, we attempted a semilog fit. The data was fit equally well by the following equation:

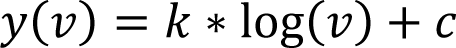

where,

*y*(*v*) = Peak firing rate, *v* = Angular velocity of flagellar movement,

*c* = Intercept, and

*k* = Slope of the line on the semilog plot.

### Multiple models explain the relationship between angular velocity and peak firing rate

For a constant change in position, the peak firing rate increased with angular velocity of the flagellum. For instance, the neuron in **Fig 3A i** increased its peak firing rate by 1.5 folds for 16-fold change in its angular velocity (**Fig 3B i**). To capture the relation between peak firing rate and angular velocity, we fit the data to 4 different models (see methods).

Various studies have employed a power law to capture the relation between neural activity and the rate of change of a physical quantity (Matheson, 1992; Ridgel *et al*., 2000; Zill, Büschges and Schmitz, 2011). Similar to these studies, in 19 out of 27 neurons (26 moths), peak firing rate and angular velocity followed a power law relationship (**Supplementary Table 1, Supplementary** Fig 2A). Conventionally, at least 2 decades of data are employed to fit a power law. Since our recordings lacked a 2-decade change in angular velocity and firing rate, we also attempted to fit other models that capture the relation between the angular velocity and the peak firing rate response.

We find that the peak firing rate changes linearly with a decade change in angular velocity (**Supplementary** Fig 2B, semilog fit). Fits with r^2^ values greater than 0.8 were considered to be good. In all, 19 out of 27 cells showed a velocity-sensitive response (r^2^ ≥ 0.8) (**Supplementary Table 1, Supplementary** Fig 2B). We observed velocity encoding in most recorded neurons albeit with different slopes of the lines. The relation is captured the best in 19 out of 27 neurons by a logistic function, with more number of fits with r^2^ ≥ 0.9 (**Fig 3B, Supplementary Table 1**). A linear fit underperforms with a good fit for only 14/26 neurons with r^2^ ≥ 0.8 (**Supplementary Table 1,** and **supplementary Fig 2C)**.

---

**Table S1:**
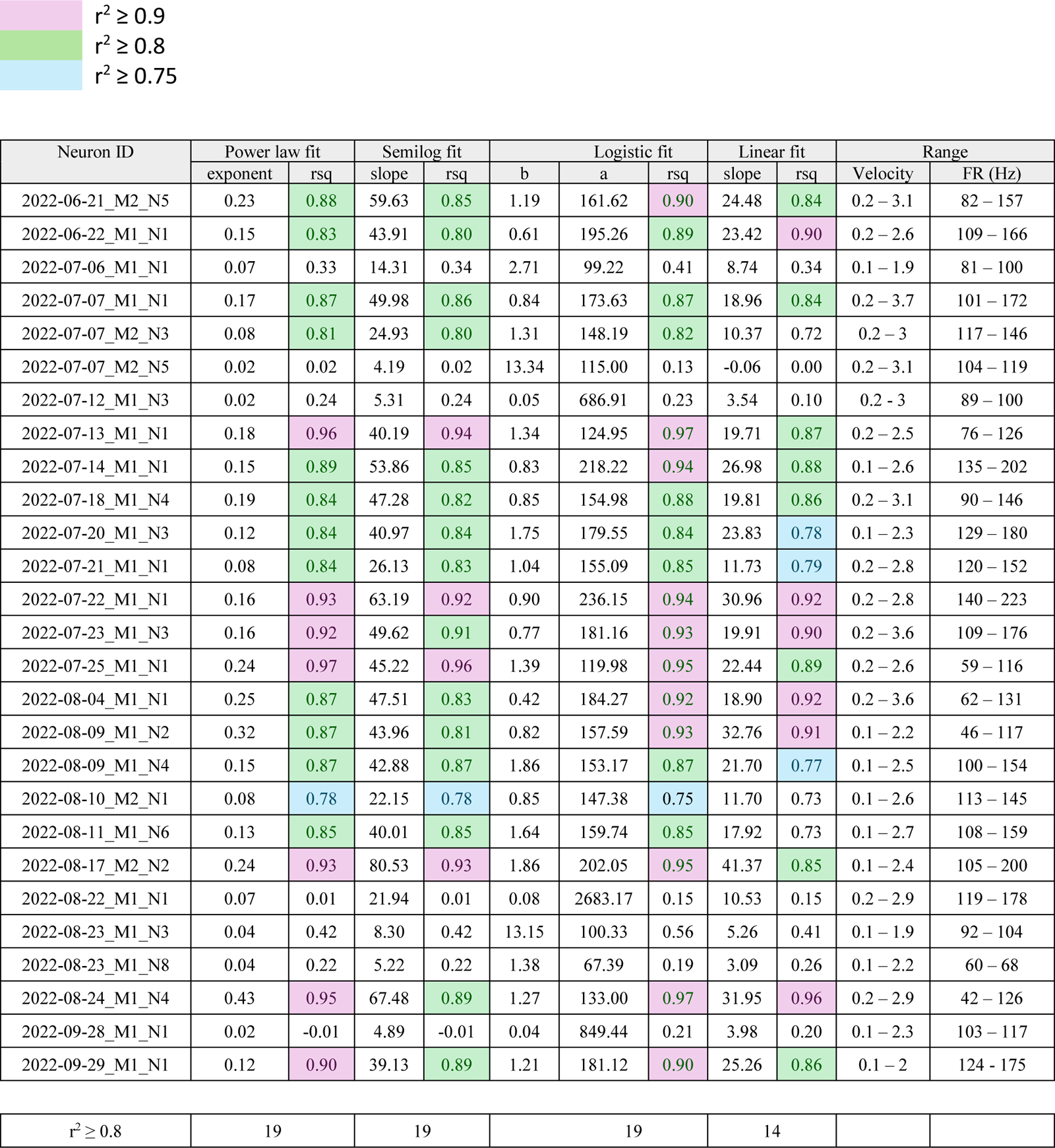
Comparison of goodness of fits for 4 models. This table compares the rsquare values of the fits of 4 models that we used to find the relation between peak firing rate and velocity. The last 2 columns display the range of angular velocities tested and the corresponding range of mean peak firing rates. The rsquare values are color-coded as follows:

**Table S2:**
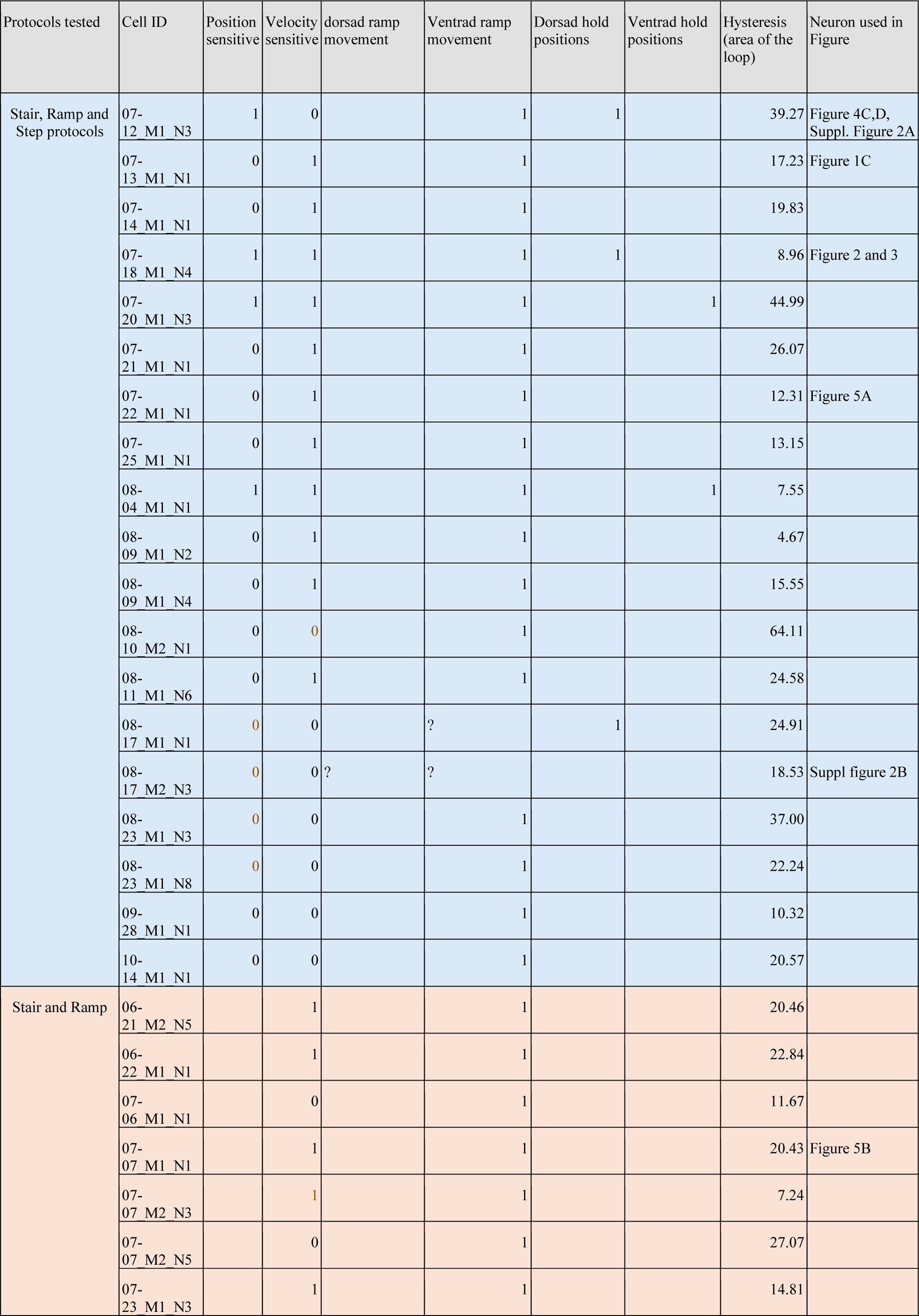
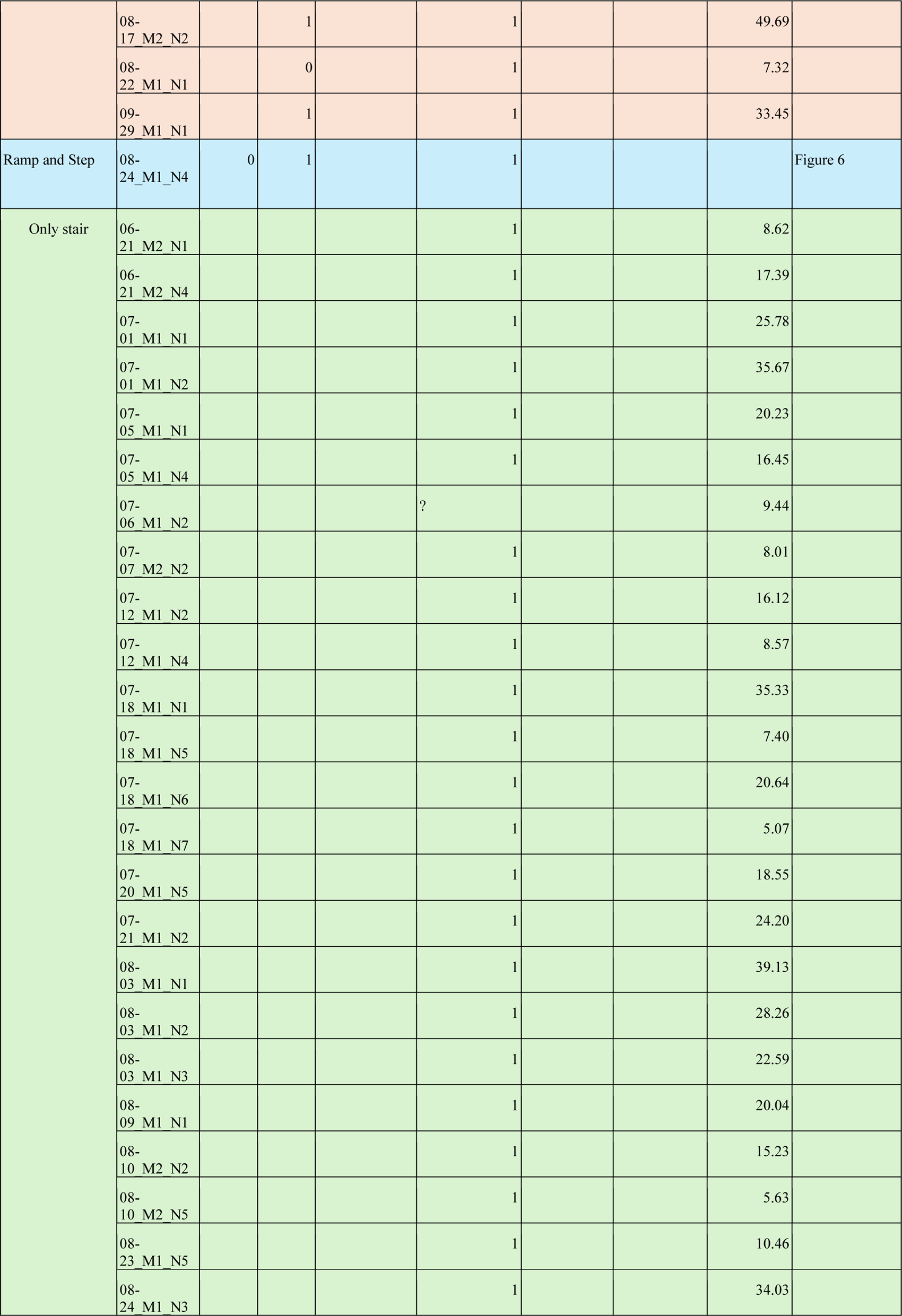
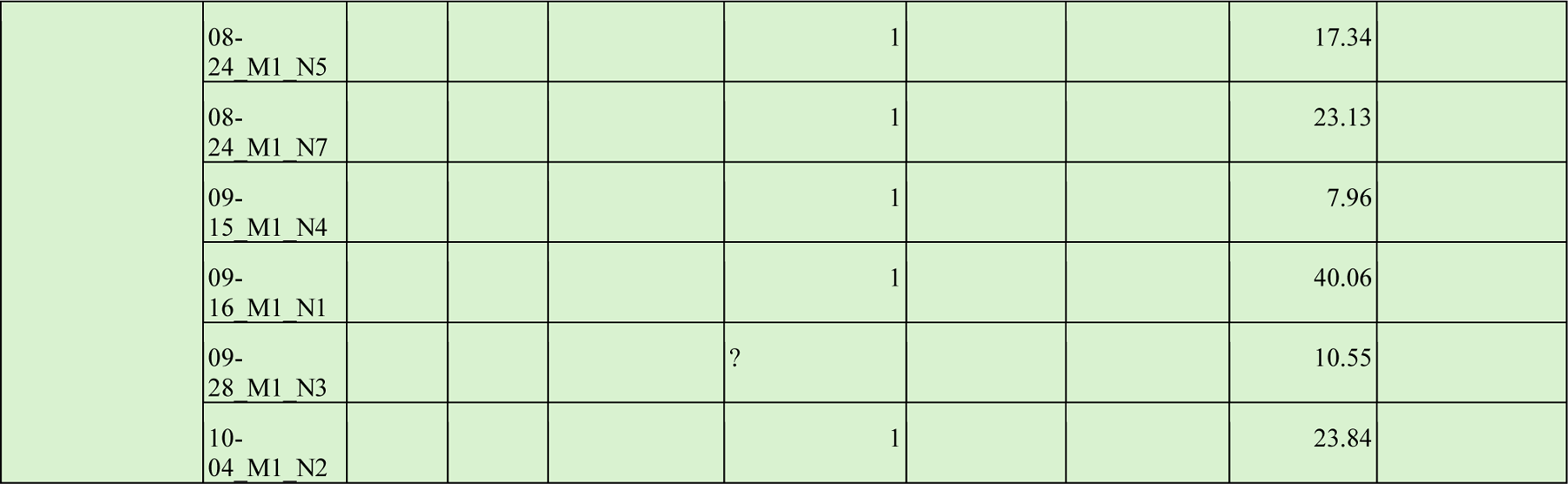
List of the neurons and their responses. Neurons used in figure are indicated in the last column of the table. In the cells, 0 : False, 1 : True,: indicates that the response was ambiguous.

## Notes

### Competing Interest Statement

The authors have declared no competing interest.

## REFERENCES

Ache, J.M. and Dürr, V. (2013) Encoding of near-range spatial information by descending interneurons in the stick insect antennal mechanosensory pathway, Journal of Neurophysiology, 110(9), pp. 2099–2112.

Adrian, E.D. and Zotterman, Y. (1926) The impulses produced by sensory nerve-endings: Part II. The response of a Single End-Organ., The Journal of physiology, 61(2), pp. 151–71.

Barth, F. G. (1964). A phasic-tonic proprioceptor in the telson of the crayfish *Procambarus clarki* (Girard). Zeitschrift für vergleichende Physiologie, 48(2), 181–189.

Barth, F.G. (2019) Mechanics to pre-process information for the fine tuning of mechanoreceptors, *Journal of Comparative Physiology A: Neuroethology, Sensory*, Neural, and Behavioral Physiology, pp. 661–686.

Benda, J. (2021) ‘Neural adaptation, Current Biology, 31(3), pp. R110–R116.

Burkhardt, D. and Gewecke, M. (1965) Mechanoreception in Arthropoda: the Chain from Stimulus to Behavioral Pattern, Cold Spring Harbor Symposia on Quantitative Biology, 30, pp. 601–614.

Burns, M.D. (1974) Structure and physiology of the locust femoral chordotonal organ., Journal of insect physiology, 20(7), pp. 1319–39.

Büschges, A. (1994). The physiology of sensory cells in the ventral scoloparium of the stick insect femoral chordotonal organ. Journal of experimental biology, 189(1), 285–292.

Bush, B.M.H. (1965) Proprioception by the Coxo-Basal Chordotonal Organ, Cb, in Legs of the Crab, Carcinus Maenas, Journal of Experimental Biology, 42(2), pp. 285–297.

Camhi, J.M. and Johnson, E.N. (1999) High-frequency steering maneuvers mediated by tactile cues: antennal wall-following in the cockroach, Journal of Experimental Biology, 202(5), pp. 631–643.

Chapman, K.M., Mosinger, J.L. and Duckrow, R.B. (1979) The role of distributed viscoelastic coupling in sensory adaptation in an insect mechanoreceptor, *Journal of Comparative Physiology ?* A, 131(1), pp. 1–12.

Chapman, K.M. and Smith, R.S. (1963) A Linear Transfer Function underlying Impulse Frequency Modulation in a Cockroach Mechanoreceptor, Nature, 197(4868), pp. 699–700.

Chatterjee, P., Prusty, A. D., Mohan, U., & Sane, S. P. (2022). Integration of visual and antennal mechanosensory feedback during head stabilization in hawkmoths. *Elife*, *11*, e78410.during head stabilization in hawkmoths., eLife, 11, pp. 1–26.

Clemens, J., Ozeri-Engelhard, N. and Murthy, M. (2018) Fast intensity adaptation enhances the encoding of sound in Drosophila, Nature Communications, 9(1), pp. 1–15.

Comer, C. and Baba, Y. (2011) Active touch in orthopteroid insects: behaviours, multisensory substrates and evolution., Philosophical transactions of the Royal Society of London. Series B, Biological sciences, 366(1581), pp. 3006–15.

Dahake, A., Stöckl, A. L., Foster, J. J., Sane, S. P., & Kelber, A. (2018). The roles of vision and antennal mechanoreception in hawkmoth flight control. Elife, 7, e37606

Dickinson, M.H. (1992) Directional Sensitivity and Mechanical Coupling Dynamics of Campaniform Sensilla During Chord-Wise Deformations of the Fly Wing, Journal of Experimental Biology, 169(1), pp. 221–233.

Dieudonne, A., Daniel, T.L. and Sane, S.P. (2014) Encoding properties of the mechanosensory neurons in the Johnston’s organ of the hawk moth, Manduca sexta, Journal of Experimental Biology, pp. 3045–3056.

Dreller, C and Kirchner, W.H. (1993) Hearing in honeybees: localization of the auditory sense organ, Journal of Comparative Physiology A, 173(3), pp. 275–279.

Dreller, C. and Kirchner, W.H. (1993) How honeybees perceive the information of the dance language, Naturwissenschaften, 80(7), pp. 319–321.

Eberl, D.F. (1999) Feeling the vibes: chordotonal mechanisms in insect hearing, Current Opinion in Neurobiology, 9(4), pp. 389–393.

Eberl, D.F., Kamikouchi, A. and Albert, J.T. (2016) Auditory Transduction, in Insect Hearing. Springer, Cham, pp. 159–175.

Field, L.H. and Matheson, T. (1998) Chordotonal Organs of Insects, in Advances in Insect Physiology, pp. 1–228.

French, A. S. (1984) Action potential adaptation in the femoral tactile spine of the cockroach,Periplaneta americana, Journal of Comparative Physiology A, 155(6), pp. 803– 812.

French, A S (1984) The receptor potential and adaptation in the cockroach tactile spine., The Journal of neuroscience 4(8), pp. 2063–8.

Gewecke, M. (1967) Die Wirkung von Luftströmung auf die Antennen und das Flugverhalten der blauen Schmeissfliege (Calliphora Erythrocephala), Zeitschrift für Vergleichende Physiologie, 54(2), pp. 121–164.

Gewecke, M. (1974) The Antennae of Insects as Air-Current Sense Organs and their Relationship to the Control of Flight, in Experimental Analysis of Insect Behaviour. Berlin, Heidelberg: Springer Berlin Heidelberg, pp. 100–113.

Göpfert, M.C. and Hennig, R.M. (2016) Hearing in Insects., Annual review of entomology, 61, pp. 257–76.

Gopfert, M.C. and Robert, D. (2001) Active auditory mechanics in mosquitoes, Proceedings of the Royal Society of London. Series B: Biological Sciences, 268(1465), pp. 333–339.

Hatsopoulos, N.G., Burrows, M. and Laurent, G. (1995) Hysteresis reduction in proprioception using presynaptic shunting inhibition, Journal of Neurophysiology, 73(3), pp. 1031–1042.

Hildebrandt, K.J., Benda, J. and Hennig, R.M. (2009) The origin of adaptation in the auditory pathway of locusts is specific to cell type and function, Journal of Neuroscience, 29(8), pp. 2626–2636.

Hofmann, T. and Bӓssler, U. (1986) Response characteristics of single trochanteral campaniform sensilla in the stick insect, Cuniculina impigra, Physiological Entomology, 11(1), pp. 17–21.

Hofmann, T. and Koch, U.T. (1985) Acceleration Receptors in the Femoral Chordotonal Organ of the Stick Insect, Cuniculina Impigra, Journal of Experimental Biology, 114(1), pp. 225–237.

Hofmann, T., Koch, U.T. and Bässler, U. (1985) Physiology of the Femoral Chordotonal Organ in the Stick Insect, Cuniculina Impigra, Journal of Experimental Biology, 114(1), pp. 207–223.

Jeram, S. and Čokl, A. (1996) Mechanoreceptors in insects: Johnstońs organ in Nezara viridula (L.) (Pentatomidae, Heteroptera), in Pflugers Archiv European Journal of Physiology. Springer Verlag, pp. R281–R282.

Johnston, C. (1855) Auditory Apparatus of the Culex Mosquito, Journal of Cell Science, S1–3(10), pp. 97–102.

Kamikouchi, A., Inagaki, H. K., Effertz, T., Hendrich, O., Fiala, A., Göpfert, M. C., & Ito, K. (2009). The neural basis of Drosophila gravity-sensing and hearing. Nature, 458(7235), 165–171.

Kladt, N. and Reiser, M.B. (2023) Drosophila antennae are dispensable for gravity orientation, bioRxiv, p. 2023.03.08.531317.

Kloppenburg, P., Camazine, S. M., Sun, X. J., Randolph, P., & Hildebrand, J. G. (1997). Organization of the antennal motor system in the sphinx moth Manduca sexta. Cell and tissue research, 287, 425–433.

Krishnan, A., Prabhakar, S., Sudarsan, S., & Sane, S. P. (2012). The neural mechanisms of antennal positioning in flying moths. Journal of Experimental Biology, 215(17), 3096–3105.

Kuster, J.E., French, A.S. and Sanders, E.J. (1983) The effects of microtubule dissociating agents on the physiology and cytology of the sensory neuron in the femoral tactile spine of the cockroach, Periplaneta americana L, Proceedings of the Royal Society of London. Series B. Biological Sciences, 219(1217), pp. 397–412.

Land, M.F. and Nilsson, D.-E. (2012) Animal Eyes. 2nd edn, Animal Eyes. 2nd edn. Oxford University Press.

Landolfa, M.A. and Miller, J.P. (1995) Stimulus-response properties of cricket cereal filiform receptors, Journal of Comparative Physiology A, 177(6), pp. 749–757.

Lapshin, D.N. and Vorontsov, D.D. (2017) Frequency organization of the Johnston organ in male mosquitoes (Diptera, Culicidae), Journal of Experimental Biology, 220(21), pp. 3927– 3938.

Lei, H., Christensen, T.A. and Hildebrand, J.G. (2004) Spatial and temporal organization of ensemble representations for different odor classes in the moth antennal lobe, Journal of Neuroscience, 24(49), pp. 11108–11119.

Loewenstein, W.R. and Mendelson, M. (1965) Components of receptor adaptation in a Pacinian corpuscle, The Journal of Physiology, 177(3), pp. 377–397.

Mamiya, A. and Dickinson, M.H. (2015) Antennal Mechanosensory Neurons Mediate Wing Motor Reflexes in Flying Drosophila, The Journal of Neuroscience, 35(20), pp. 7977–7991.

Massarelli, N., Yau, A. L., Hoffman, K. A., Kiemel, T., & Tytell, E. D. (2017). Characterization of the encoding properties of intraspinal mechanosensory neurons in the lamprey. Journal of Comparative Physiology A, 203, 831–841.

Matheson, T. (1992) Range fractionation in the locust metathoracic femoral chordotonal organ, Journal of Comparative Physiology A, 170(4), pp. 509–520.

Nakajima, S. and Onodera, K. (1969a) Adaptation of the generator potential in the crayfish stretch receptors under constant length and constant tension, The Journal of Physiology, 200(1), pp. 187–204.

Nakajima, S. and Onodera, K. (1969b) Membrane properties of the stretch receptor neurones of crayfish with particular reference to mechanisms of sensory adaptation, The Journal of Physiology, 200(1), pp. 161–185.

Natesan, D., Saxena, N., Ekeberg, Ö., & Sane, S. P. (2019). Tuneable reflexes control antennal positioning in flying hawkmoths. Nature communications, 10(1), 5593..

Niehaus, M. (1981) Flight and flight control by the antennae in the Small Tortoiseshell (*Aglais urticae L.*, Lepidoptera), *Journal of Comparative Physiology ?* A, 145(2), pp. 257– 264.

Okada, J. and Toh, Y. (2000) The role of antennal hair plates in object-guided tactile orientation of the cockroach ( Periplaneta americana ), *Journal of Comparative Physiology A: Sensory*, Neural, and Behavioral Physiology, 186(9), pp. 849–857.

Patella, P. and Wilson, R.I. (2018) Functional Maps of Mechanosensory Features in the Drosophila Brain, Current Biology, 28(8), pp. 1189–1203.e5.

Ridgel, A. L., Frazier, S. F., DiCaprio, R. A., & Zill, S. N. (2000). Encoding of forces by cockroach tibial campaniform sensilla: implications in dynamic control of posture and locomotion. Journal of Comparative Physiology A, 186, 359–374.

Khurana, T. and Sane, S.P. (2016) Airflow and optic flow mediate antennal positioning in flying honeybees, eLife, 5.

Sane, S. P., Dieudonné, A., Willis, M. A., & Daniel, T. L. (2007). Antennal mechanosensors mediate flight control in moths. science, 315(5813), 863–866.

Sane, S.P., Manjunath, M. and Mukunda, C.L. (2023) Vestibular feedback for flight control in hawkmoths., Trends in neurosciences, 46(8), pp. 614–616.

Sane, S.P. and McHenry, M.J. (2009) The biomechanics of sensory organs, Integrative and Comparative Biology, 49(6), pp. i8–i23.

Sant, H.H. and Sane, S.P. (2018) The mechanosensory-motor apparatus of antennae in the Oleander hawk moth (Daphnis nerii, Lepidoptera)., The Journal of comparative neurology, 526(14), pp. 2215–2230.

Schneider, D. (1964) Insect Antennae, Annual Review of Entomology, 9(1), pp. 103–122.

Shimozawa, T. and Kanou, M. (1984) The aerodynamics and sensory physiology of range fractionation in the cereal filiform sensilla of the cricket Gryllus bimaculatus, Journal of Comparative Physiology A, 155(4), pp. 495–505.

Simon, M.A. and Trimmer, B.A. (2009) Movement encoding by a stretch receptor in the soft-bodied caterpillar, Manduca sexta, Journal of Experimental Biology, 212(7), pp. 1021–1031.

Suver, M. P., Matheson, A. M., Sarkar, S., Damiata, M., Schoppik, D., & Nagel, K. I. (2019). Encoding of wind direction by central neurons in Drosophila. Neuron, 102(4), 828–842.

Thorson, J. and Biederman-Thorson, M. (1974) Distributed Relaxation Processes in Sensory Adaptation, Science, 183(4121), pp. 161–172.

Yorozu, S., Wong, A., Fischer, B. J., Dankert, H., Kernan, M. J., Kamikouchi, A., … & Anderson, D. J. (2009). Distinct sensory representations of wind and near-field sound in the Drosophila brain. Nature, 458(7235), 201–205.

Zanini, D. and Göpfert, M.C. (2014) TRPs in hearing, Handbook of Experimental Pharmacology, 223, pp. 899–916.

Zill, S.N. (1985) Plasticity and proprioception in insects. I. Responses and cellular properties of individual receptors of the locust metathoracic femoral chordotonal organ., The Journal of experimental biology, 116(1), pp. 435–61.

Zill, S.N., Büschges, A. and Schmitz, J. (2011) Encoding of force increases and decreases by tibial campaniform sensilla in the stick insect, Carausius morosus, Journal of Comparative Physiology A, 197(8), pp. 851–867.

Zill, S.N., Moran, D.T. and Varela, F.G. (1981) The exoskeleton and insect proprioception ii. Reflex effects of tibial campaniform sensilla in the american cockroach, *Periplaneta americana*, J. exp. Biol, 94, pp. 43–55.

